# Far-red chemigenetic biosensors for multi-dimensional and super-resolved kinase activity imaging

**DOI:** 10.1101/2024.02.10.579766

**Authors:** Michelle S. Frei, Samantha A. Sanchez, Longwei Liu, Falk Schneider, Zichen Wang, Hiroyuki Hakozaki, Yajuan Li, Anne C. Lyons, Theresa V. Rohm, Jerrold M. Olefsky, Lingyan Shi, Johannes Schöneberg, Scott E. Fraser, Sohum Mehta, Yingxiao Wang, Jin Zhang

**Affiliations:** Department of Pharmacology, University of California, San Diego, La Jolla, CA, USA; Shu Chien-Gene Lay Department of Bioengineering, University of California, San Diego, La Jolla, CA, USA; Institute of Engineering in Medicine, University of California, San Diego, La Jolla, CA, USA; Alfred E. Mann Department of Biomedical Engineering, University of Southern California, Los Angeles, CA, USA; Translational Imaging Center, University of Southern California, Los Angeles, CA, USA; Dana and David Dornsife College of Letters, Arts and Sciences, University of Southern California, Los Angeles, CA, USA; Department of Chemistry and Biochemistry, University of California, San Diego, La Jolla, CA, USA; Division of Endocrinology and Metabolism, Department of Medicine, University of California, San Diego, La Jolla, CA, USA; Department of Biological Sciences, Division of Molecular and Computational Biology, University of Southern California, Los Angeles, CA, USA

## Abstract

Fluorescent biosensors revolutionized biomedical science by enabling the direct measurement of signaling activities in living cells, yet the current technology is limited in resolution and dimensionality. Here, we introduce highly sensitive chemigenetic kinase activity biosensors that combine the genetically encodable self-labeling protein tag HaloTag7 with bright far-red-emitting synthetic fluorophores. This technology enables five-color biosensor multiplexing, 4D activity imaging, and functional super-resolution imaging via stimulated emission depletion (STED) microscopy.

## Introduction

Genetically encoded biosensors are powerful tools to investigate the dynamic biochemical processes underlying physiological functions and pathological derailment^1^. Despite numerous advances, biosensor technology continues to lag in three major areas. First, sensitive and bright signaling biosensors remain limited to the cyan-to-green spectral region^2–5^, drastically restricting our capability to monitor multiple biochemical and signaling activities simultaneously. Secondly, there is an urgent need for fluorescent biosensors capable of withstanding the photophysical demands of tissue-scale 4D activity imaging to track biochemical processes in real time within physiologically relevant contexts. Finally, new biosensor designs are needed to make functional super-resolution microscopy^6–8^ a practical reality for investigating the compartmentalization of signaling networks beyond the diffraction limit and mapping nanoscale biochemical activity architecture^9,10^.

## Results and Discussion

Recognizing that chemigenetic fluorescent biosensors using synthetic fluorophores and self-labeling protein tags^11–14^ could offer spectral versatility into the far-red, high photostability, and high signal-to-background ratios, we strived to develop chemigenetic far-red kinase activity reporters to address the aforementioned technological gaps. Specifically, we combined a sensing unit consisting of a protein kinase A (PKA)-specific substrate peptide and phosphoamino acid-binding (PAAB) forkhead-associated 1 (FHA1) domain^2,15^ with a circularly permutated HaloTag reporting unit (Figure 1a, Supplementary Figure S1)^12,14^, which we labeled with the fluorogenic silicon-rhodamine JF_635_-chloroalkane (JF_635_-CA)^16^. Analogous to recent chemigenetic Ca^2+^ biosensors^12,14^, we hypothesized that the kinase-activity-induced conformational change caused by binding of the phosphorylated substrate to FHA1 would influence the open-close equilibrium between the colorless, non-fluorescent spirolactone and the colored, fluorescent zwitterion, and thus the fluorescence intensity of JF_635_ (Figure 1a-b). We optimized the order of protein domains and the lengths of the interdomain linkers through four rounds of rational engineering. The best candidate, **Halo**Tag-based **A k**inase **a**ctivity **r**eporter 1.0 (HaloAKAR1.0-JF_635_), showed good dynamic range (ΔF/F_0_ = 129.8%, Supplementary Table S1-2) in living HeLa cells upon maximal PKA stimulation with the adenylate cyclase activator forskolin (Fsk) and the phosphodiesterase inhibitor 3-isobutyl-1-methylxanthine (IBMX). However, the biosensor was rather dim in its basal state (JF_635_/EGFP = 0.27). We therefore set out to improve the biosensor’s dynamic range and basal brightness simultaneously.

**Fig. 1:**
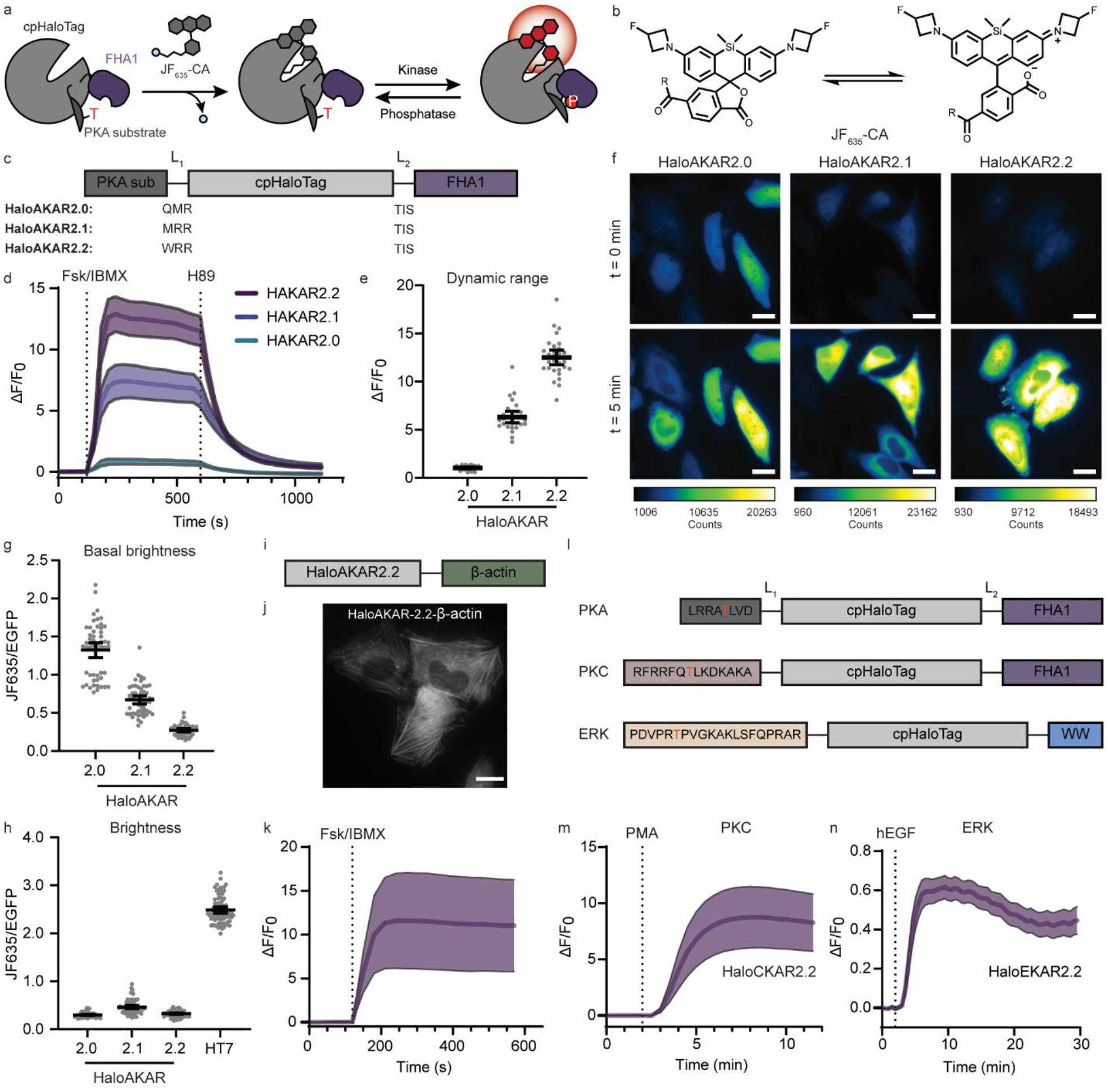
Design and characterization of HaloAKARs. **a**, Schematic of chemigenetic HaloAKAR biosensor based on cpHaloTag labeled with JF_635_-CA and a phosphorylation-dependent switch that influences JF_635_ fluorescence intensity. **b**, Open-close equilibrium of JF_635_-CA, R = chloroalkane (CA). **c**, Domain structure of the three selected HaloAKARs, including linker sequences. PKA sub: PKA substrate. **d**, Average time courses of HeLa cells expressing HaloAKAR2.0, HaloAKAR2.1, or HaloAKAR2.2 labeled with JF_635_-CA and stimulated with 50 μM Fsk/100 μM IBMX and 20 μM H89 (n = 7, 10, and 13 cells for 2.0, 2.1, and 2.2, respectively). **e**, Maximum Fsk/IBMX-stimulated response (ΔF/F_0_) for HaloAKAR-JF_635_ biosensors. **f**, Representative pseudo-color widefield images of HeLa cells expressing HaloAKARs labeled with JF_635_-CA before and after Fsk/IBMX stimulation. Scale bars, 20 μm. **g-h**, Basal (**g**) and activated brightness (**h**) of HaloAKAR-JF_635_ biosensors. In h, the brightness of HaloTag7-JF_635_ is given for comparison. **i**, Domain structure of β-actin-targeted HaloAKAR2.2. **j**, Representative image of HeLa cells expressing HaloAKAR2.2-β-actin after Fsk/IBMX stimulation. Scale bar, 20 μm. **k**, Average time course of HeLa cells expressing HaloAKAR2.2-β-actin labeled with JF_635_-CA upon Fsk/IBMX stimulation (n = 4 cells). **l**, Domain structure of HaloTag-based PKA, PKC and ERK biosensors (HaloAKAR, HaloCKAR, and HaloEKAR). **m**-**n**, Average time courses of HeLa and HEK293T cells expressing HaloCKAR2.2 (**m**) and HaloEKAR2.2 (**n**) labeled with JF_635_-CA stimulated with phorbol 12-myristate 13-acetate (PMA, 100 ng mL^-1^) or epidermal growth factor (EGF, 100 ng mL^-1^), respectively (n = 9 and 35 cells). Time courses are representatives of three replicates, and dashed lines indicate addition of drug. Solid lines indicate mean responses; shaded areas correspond to 95% confidence interval. For average measurements, individual data points are shown from three independent repeats along with mean and 95% confidence intervals. Numbers of cells (n) can be found in Supplementary Table S5.

Using a Sort-Seq^17,18^ approach, we screened >15,000 biosensor variants and identified three new HaloAKAR variants (Figure 1c-h, Supplementary Figure S2, Supplementary Table S3-5). HaloAKAR2.0-JF_635_ had high basal brightness and good dynamic range (ΔF/F_0_ = 104.4%, JF_635_/EGFP = 1.32), although still dimmer than HaloTag7-JF_635_. HaloAKAR2.1-JF_635_ showed good basal brightness and high dynamic range (ΔF/F_0_ = 634%, JF_635_/EGFP = 0.67), and HaloAKAR2.2-JF_635_ exhibited low basal brightness but extremely high dynamic range (ΔF/F_0_ = 1250%, JF_635_/EGFP = 0.27). All HaloAKARs showed a reversible response upon treatment with the PKA inhibitor H89 and showed no intensity change when the phospho-acceptor Thr on the substrate peptide was mutated to Ala, nor upon stimulation of related ACG kinases (Supplementary Figure S3), indicating that their responses were specific to phosphorylation by PKA.

Purified HaloAKARs rapidly reacted with fluorophore substrate *in vitro*, albeit at a slower rate than HaloTag7 (Supplementary Figure S4a-d, Supplementary Table S6). Changes in HaloAKARs-JF_635_ fluorescence emission (ΔF/F_0_ = 45, 291, and 746% for 2.0, 2.1, and 2.2, respectively) mostly stem from changes in extinction coefficient ε (Δε/ε_0_ = 42, 165, and 328%), whereas quantum yield ϕ has no (HaloAKAR2.0: Δϕ/ϕ_0_ = 2%) or small influence (HaloAKAR2.1 and 2.2: Δϕ/ϕ_0_ = 48 and 98%). Additionally, we found that the fluorescence lifetime of HaloAKARs-JF_635_ changed only marginally upon PKA activity (Supplementary Figure S4e-g, Supplementary Table S7). Our data strengthen the model whereby biosensor fluorescence intensity increases are based on a shift in the open-close equilibrium, leading to a larger fraction of fluorophore in the absorbing state and hence an increase in apparent ε. This is further supported by the responses of HaloAKARs labeled with fluorophores whose equilibria are shifted more towards the zwitterionic form compared to JF_635_ (JF_639_, JFX_646_ and JF_669_), which showed smaller changes upon Fsk/IBMX stimulation than fluorophores with comparable equilibria (JF_585_, Supplementary Figure S5). The use of JF_585_ additionally expands the utility of this platform, giving access to high-performance red biosensors.

HaloAKAR2.2-JF_635_ shows the highest dynamic range reported to date for any kinase biosensor, surpassing ExRai-AKAR2 (480/405 excitation ratio ΔR/R_0_ = 1,095%)^2^, which is in the green spectral region, and drastically outperforming the few available NIR Förster resonance energy transfer (FRET) biosensors (< 40%)^4,5^. This allowed for sensitive measurements of PKA activity at low doses of Fsk (Supplementary Figure S6). Benefiting from their high dynamic ranges, HaloAKARs also enable sensitive PKA activity measurements at different subcellular locations, including the plasma membrane, clathrin-coated pits, the actin or microtubule network, or the outer mitochondrial membrane (Figure 1i-k, Supplementary Figure S7). Further, this chemigenetic biosensor design can be expanded to other kinases by switching the PKA-specific peptide for peptides specific to other kinases, such as protein kinase C (PKC) or protein kinase B (Akt, Figure 1l-m). By further replacing the FHA1 domain with a WW domain, we generated biosensors for extracellular signal-regulated kinase (ERK). Replacing the PAAB domain had a larger impact on the dynamic range than switching out the substrate. Nevertheless, HaloEKARs showed dynamic ranges as high as 60% (HaloEKAR2.2, Figure 1n, Supplementary Figure S8). Taken together, HaloTag-based KARs represent an ideal platform to generate sensitive and specific far-red- and red-emitting biosensors for diverse kinases with promising photophysical properties for advanced bioimaging applications.

These far-red chemigenetic biosensors are ideally suited for multiplexed activity sensing with fluorescent protein-based biosensors in other spectral regions. We first performed three-color imaging using HaloAKAR2.2-JF_635_, the red Ca^2+^ biosensor RCaMP1d^19^, and the green cAMP biosensor GFlamp-1^20^ in MIN6 β-cells. Stimulation with the K^+^ channel blocker tetraethylammonium chloride led to in-phase oscillations of Ca^2+^ and PKA activity and inverse oscillations for cAMP (Figure 2a, Supplementary Figure S9), providing single-cell capture of the previously identified oscillatory circuit^21^ with unprecedented clarity. Further combination with sapphire and blue biosensors allowed for four- and five-color imaging (Supplementary Figures S10-11). For instance, combining HaloAKAR2.1-JF_635_ with the red cAMP biosensor pinkFlamindo, the yellow cGMP biosensor cGull^22^, the sapphire PKC biosensor sapphireCKAR^3^, and the blue Ca^2+^ biosensor BGeco^23^ allowed us to follow the concentration and activities of all five analytes upon pharmacological stimulation (Figure 2b). This record-high five-color biosensor multiplexing in the same living cell not only underscores the strengths of this technology, but also sets a new benchmark for future advancements in functional live-cell microscopy.

**Figure 2:**
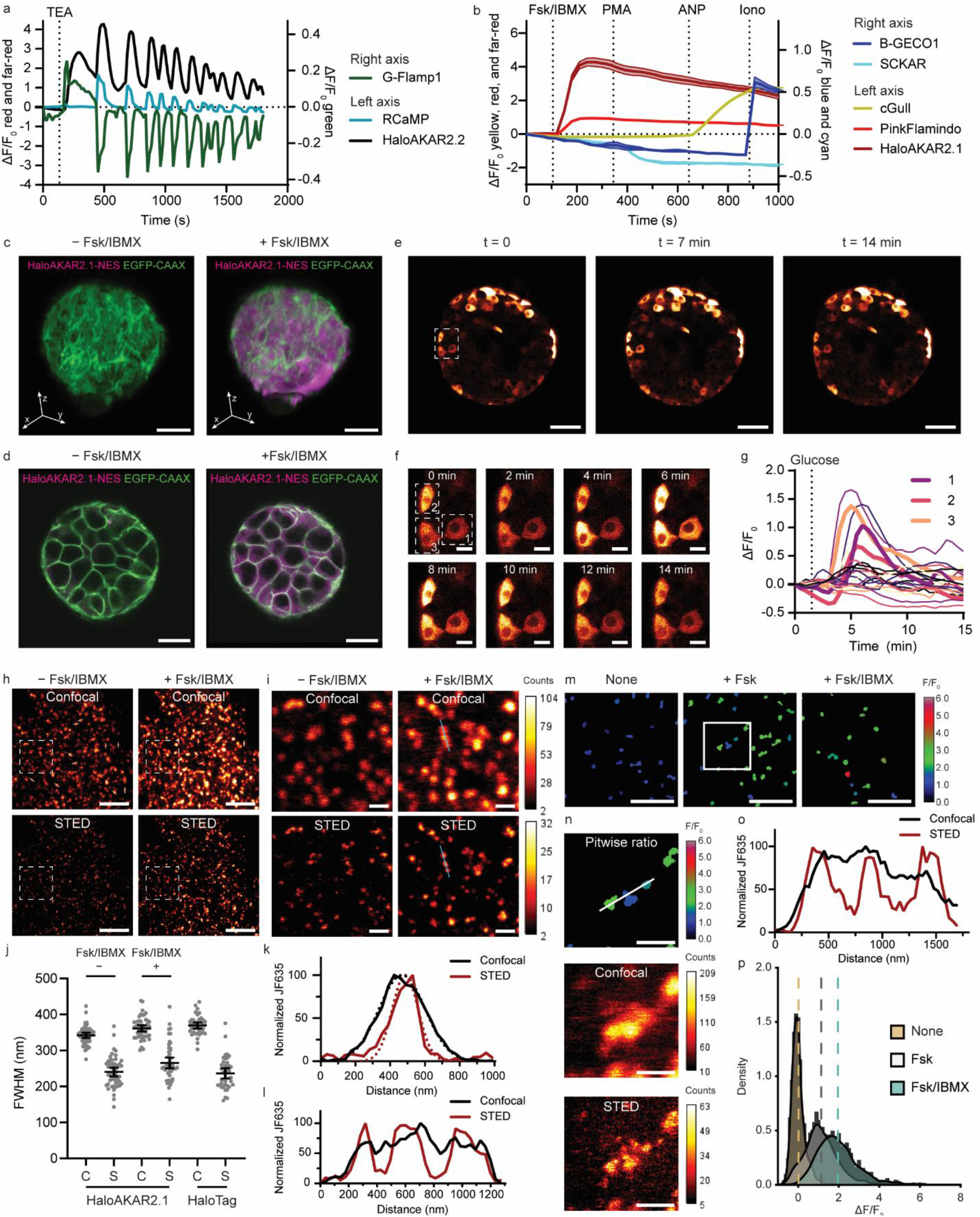
Application of HaloAKARs in multiplexed biosensor imaging, 4D activity imaging and super-resolution microscopy. **a** Three-color multiplexing of cAMP (G-Flamp1, green), Ca^2+^ (RCaMP, blue) and PKA activity (HaloAKAR2.2, black) in single MIN6 cells stimulated with 20 mM tetraethylammonium chloride (TEA). **b**, Five-color multiplexing of Ca^2+^ (B-GECO1, blue), PKC activity (sapphireCKAR (SCKAR), cyan), cGMP (cGull, yellow), cAMP (pinkFlamindo, red), and PKA activity (HaloAKAR2.1, dark-red) in HeLa cells stimulated with 50 μM Fsk/100 μM IBMX (Fsk/IBMX), 100 ng mL^-1^ PMA, 0.4 μM atrial natriuretic peptide (ANP), and 1 μM ionomycin (Iono). Solid lines indicate mean response; shaded areas correspond to SD (n = 2 cells). **c-d**, Imaging of 4-d-old HEK293T spheroids stably expressing cytosolic HaloAKAR2.1 (HaloAKAR2.1-NES, magenta) and plasma membrane anchored EGFP (EGFP-CAAX, green) labeled with JF_635_-CA via lattice light-sheet microscopy (**c**) and 2P microscopy (**d**). Representative 3D renderings (**c**) and overlay images of one z-plane (**d**) before and after Fsk/IBMX stimulation are shown. Scale bars, 20 μm. **e-g**, Confocal imaging of isolated pancreatic islets expressing HaloAKAR2.1-T2A-EGFP upon stimulation with 25 mM glucose. Overview (**e**) and zoomed in images (**f**) are shown, along with single-cell traces (**g**) highlighting the three cells in f. **h-i**, Representative confocal and STED images of HeLa cells expressing HaloAKAR2.1-clathrin-light-chain (CLC) labeled with JF_635_-CA before and after Fsk/IBMX stimulation. Overview (**h**) and zoom (**i**) images are shown. Lines used to quantify the diameter of clathrin-coated pits are indicated in the confocal images (h, white), as well as the line through three neighboring pits (i, blue). Scale bars, 5 μm (**h**) and 1 μm (**i**). **j**, Diameter of clathrin-coated pits. Quantifications are given for HaloAKAR2.1-CLC before and after Fsk/IBMX stimulation, as well as HaloTag7-CLC, for both confocal (C) and STED (S) images. Individual data points are shown along with mean and 95% confidence interval. HaloAKAR2.1, n = 50 pits from 5 cells from 5 replicates; HaloTag7, n = 40 pits from 4 cells from 4 replicates. **k-l**, Representative normalized line profiles (solid) through a clathrin-coated pit for diameter quantification (**k**) or three neighboring pits (**l**) resolved in STED. Confocal (black) and STED (red) curves are shown, as well as fitted curves (dotted) in k. **m**, Pitwise activity analysis showing the mean F/F_0_ for pits present after stimulation. Representative images shown under three conditions: untreated (left), 50 μM Fsk (middle), and 50 μM Fsk/100 μM (right). Scale bars, 2.5 μm. **n**, Zoom of outline region in (**m**) showing pitwise ratio (top), as well as confocal (middle) and STED (bottom) images after Fsk stimulation. Scale bars, 500 nm. **o**, Representative normalized line profile through the three clathrin-coated pits, revealing that they are resolved in STED but not in confocal microscopy. **p**, Density plots of mean ΔF/F_0_ per clathrin-coated pit. The overall mean is given as a dashed line (no treatment: 900 pits from 5 cells from 5 replicates; Fsk: 934 pits from 7 cells from 7 replicates; Fsk/IBMX: 1377 pits from 9 cells from 9 replicates).

Motivated by the far-red spectral characteristics of HaloAKARs-JF_635_ and their favorable bleaching kinetics compared with the chemigenetic Ca^2+^ biosensor HaloCaMP1a-JF_635_ (Supplementary Figure S4h-j, Supplementary Table S8), we tested their application for 4D activity imaging in more physiologically relevant 3D tissues. First, we imaged 4-d-old HEK293T spheroids (diameter ∼60 μm) stably expressing HaloAKAR2.1-NES and EGFP-CAAX via spinning disc confocal and lattice light-sheet microscopy (Figure 2c-d, Supplementary Figures S12-13). PKA responses were visible upon Fsk/IBMX treatment in both modalities, clearly demonstrating the ability of HaloAKAR2.1-JF_635_ to enable 4D activity imaging. Nonetheless, residual bleaching is still visible and further optimization is required. Secondly, two-photon (2P) microscopy allowed imaging of HaloAKAR2.1-JF_635_ and EGFP using a single laser line (1031 nm), reducing the excitation light needed (Supplementary Figure S14). Moreover, HaloAKAR2.1 could be labeled with JF_585_-CA and excited at 1031 nm, leading to a stronger signal, as expected from this fluorophore’s 2P excitation spectrum^16^. Hence, both red and far-red HaloAKARs allow imaging via 2P microscopy. Lastly, we expressed HaloAKAR2.1 in isolated pancreatic islets via AAV infection. Upon glucose stimulation, confocal microscopy showed asynchronous PKA activity fluctuations in individual β-cells (Figure 2e-g, Supplementary Figure S15 and Supplementary Video S1). While islet Ca^2+^ imaging has been performed for many years^24^, the PKA activity imaging in pancreatic islets demonstrated here opens the door for direct interrogation of critical signaling changes in response to important G-protein-coupled receptor ligands, such as glucagon-like peptide 1 receptor agonists^25,26^. These results demonstrate the power of far-red HaloAKARs for 4D activity imaging in physiologically relevant systems.

The high brightness and photostability of synthetic fluorophores make them ideally suited for live-cell super-resolution microscopy^27,28^. To test HaloAKAR performance, we used STED microscopy to image PKA activity in HeLa cells expressing clathrin-targeted HaloAKAR2.1 (Figure 2h-l). Analysis of the average diameter of clathrin-coated pits in confocal and STED images revealed a clear resolution improvement both before (FWHM_C_ = 342±4 nm; FWHM_S_ = 241±6 nm) and after Fsk/IBMX treatment (FWHM_C_ = 362±4 nm; FWHM_S_ = 266±7 nm), which was comparable to HaloTag7-labeled clathrin-coated pits (FHWM_C_ = 370±4 nm; FHWM_S_ = 237±7 nm, Supplementary Figure S16). Pit-wise analysis of ΔF/F_0_ revealed higher mean activity following treatment with Fsk/IBMX (1.9±0.6) versus Fsk alone (1.1±0.6). The broad activity distributions seen with both treatments indicate heterogeneous activity levels on the single-pit level. Most importantly, the enhanced spatial resolution of STED microscopy allowed us to measure PKA activity at individual pits that were not resolvable under confocal microscopy, allowing better capture of signaling heterogeneity (Figure 2m-p, Supplementary Figure S17). HaloAKAR therefore paves the way for future studies of compartmentalized PKA activity at the nanoscale.

In summary, HaloAKARs are a series of chemigenetic PKA activity biosensors with red or far-red spectral properties and exceptionally high dynamic ranges. HaloAKARs expanded biosensor multiplexing, enabled 4D activity imaging, and realized STED-based super-resolution activity imaging. We expect this technology will lead the way in illuminating signaling activity dynamics across scales.

## Data Availability

Plasmids encoding pcDNA3.1-HaloAKAR2.0-T2A-EGFP, pcDNA3.1-HaloAKAR2.1-T2A-EGFP, pcDNA3.1-HaloAKAR2.2-T2A-EGFP, pcDNA3.1-HaloAKAR2.1-NES-T2A-EGFP-CAAX, pcDNA3.1-HaloAKAR2.1-CLC, pcDNA3.1-HaloCKAR2.2-T2A-EGFP, pcDNA3.1-HaloAktKAR2.2-T2A-EGFP, pcDNA3.1-HaloEKAR2.2-T2A-EGFP will be deposited on Addgene. The stable cell line generated is available upon request. Source data are provided with this paper. Further data supporting the findings of this study are available upon reasonable request.

## Code Availability

Custom MATLAB scripts for clathrin-coated pit analysis will be available at https://github.com/jinzhanglab-ucsd. Custom ImageJ macros, R code, and Python code used to analyze imaging data, as well as bash and R scripts used to analyze sequencing data, are available upon reasonable request.

## Acknowledgements

We thank L. Lavis and J. Grimm (Janelia Research Campus) for providing JF fluorophores. We thank I. Garcia (USC) for assistance with cell culture. We thank C. Alvarez (UCSD) for assistance with cloning. We thank J. Bogomolovas and J. Chen (UCSD) for access and assistance to their fluorescence polarization plate reader. We thank J. Chung and J. Yang (UCSD) for access and assistance to their fluorescence and absorbance spectrometer. We thank Z. Liang (UCSD) for help with processing of sequencing results. We thank H. Farrants (Janelia Research Campus) for discussions on chemigenetic biosensors. We thank A. Linnemann (Indiana University) for providingthe pAAV-Ins-GCaMP plasmid map. We thank Q. Ni (UCSD) for support in material acquisition and tissue culture. We thank P. Guo from the UC San Diego Nikon Imaging Center, as well as J. Santini and M. Erb from the UC San Diego School of Medicine Microscopy Core, for assistance with microscopy. We thank the Translational Imaging Center, A. Shwartz and J. Junge, at USC for their help with STED microscopy. This publication includes data generated at the UC San Diego IGM Genomics Center utilizing an Illumina NovaSeq 6000 that was purchased with funding from a National Institutes of Health SIG grant (#S10 OD026929). The Leica SP8 in the UC San Diego School of Medicine Microscopy Core was funded by NINDS P30NS04710.

## Author Contributions

M.S.F. and J.Z. conceived and designed the study. M.S.F. performed the rational and Sort-Seq screen; performed in cellulo and in vitro characterization of all HaloAKAR variants; performed widefield, widefield multiplexing, spinning disc confocal, scanning confocal, FLIM and STED microscopy and analysis thereof; S.A.S. generated and characterized HaloKAR variants; generated spheroids and performed spinning disc confocal microscopy and analysis thereof; M.S.F. and S.A.S. generated stable cell lines, assisted 2P and lattice light-sheet microscopy and performed the analysis of the former, and performed islets experiments by scanning confocal microscopy; L.L. assisted the Sort-Seq screen and performed FACS; F.S. assisted STED microscopy; Z.W. processed and analyzed lattice light-sheet data; H.H. performed lattice light-sheet microscopy; Y.L. performed 2P microscopy; A.C.L. generated MATLAB scripts for object identification and pairing in STED images; T.V.R. isolated mouse islets; M.S.F., J.M.O., L.S., J.S., S.E.F., S.M., Y.W. and J.Z. supervised the work; M.S.F., S.M. and J.Z. wrote the manuscript with input from all authors.

## Funding

This research was supported by the Swiss National Science Foundation (SNSF) grants P2ELP3_199834 and P500PN_214234 to M.S.F.; the Charles Lee Powell Foundation Powell Fellowship at the University of California, San Diego (UCSD) and the UCSD Interfaces Graduate Training Program National Institutes of Health (NIH)/National Institute of Biomedical Imaging and Bioengineering (NIBIB) T32 Training in Multi-scale Analysis of Biological Structures and Function Training Grant 5T32EB009380-15 to S.A.S.; an EMBO (ALTF 849-2020) and HFSP (LT000404/2021-L) fellowship to F.S.; the National Science Foundation (NSF) Graduate Research Fellowship (DGE-2038238) and American Heart Association Predoctoral Fellowship (24PRE1186687) to A.C.L. Any opinions, findings, and conclusions or recommendations expressed in this material are those of the author(s) and do not necessarily reflect the views of the NSF; the Swiss National Science Foundation (P2BSP3_200177) and the Larry L. Hillblom Foundation (2023-D-012-FEL) to T.V.R.; the U.S. National Institute of Diabetes and Digestive and Kidney Diseases (DK063491 and DK101395) and a grant from Janssen Pharmaceuticals to J.M.O.; NIH R01GM149976, NIH U01AI167892, NIH 5R01NS111039, NIH R21NS125395, NIHU54DK134301, NIHU54 HL165443, and UCSD Startup funds to L.S.; NIH grant R01 CA262815 to J.Z and Y.W.; NIH grants EB029122, R35 GM140929, R01 HL121365, and HD107206 to Y.W.; NIH grants R35 CA197622, R01 DK073368, R01 DE030497, R01 HL162302 and RF1 MH126707 to J.Z.

## Competing Interests

We declare that none of the authors have competing financial or non-financial interests.

## Notes

### Competing Interest Statement

The authors have declared no competing interest.

