## Supplementary Information for "Far-red chemigenetic biosensors for multi-dimensional and super-resolved kinase activity imaging"

### Supplementary Figures

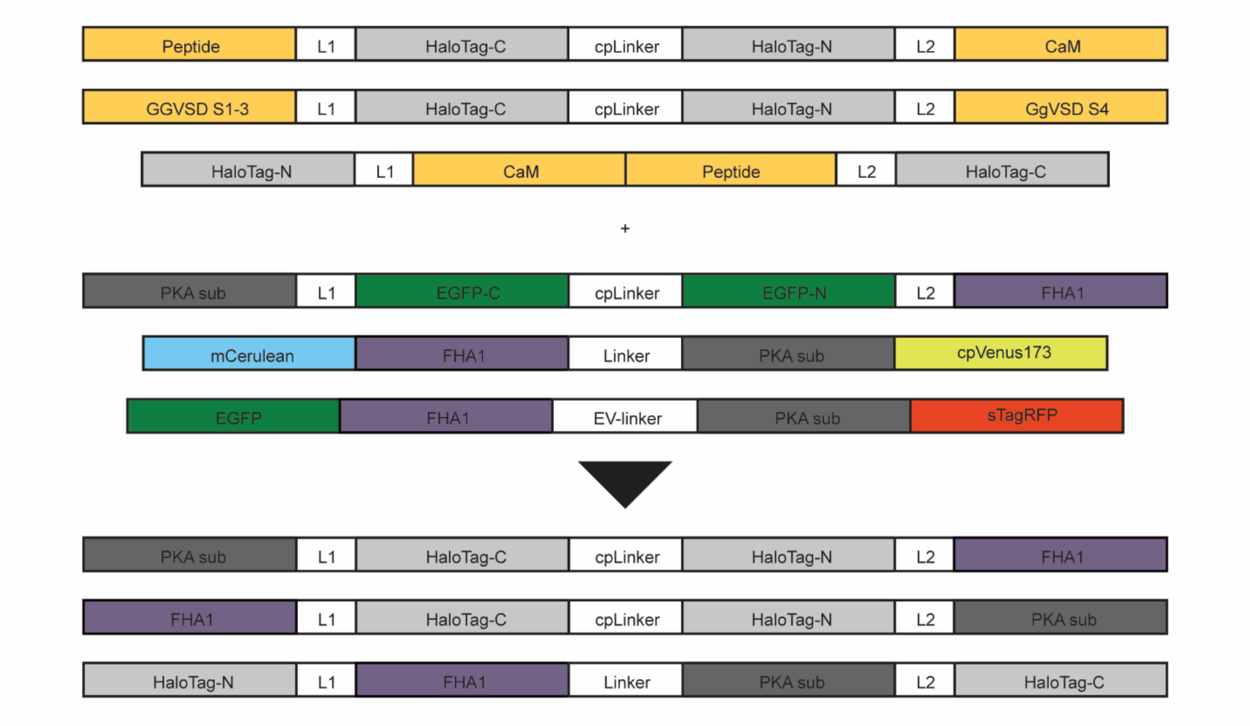

**Supplementary Figure S1**: Domain structures of biosensors generated in this work. Chemigenetic biosensors based on cpHaloTag[1] and HaloTag[2] for calcium and voltage were used as templates and combined with the sensing units derived from ExRaiAKAR1[3] or ExRaiAKAR2[4] or the FRET biosensors AKAR4[5] or GR-AKAR[6].

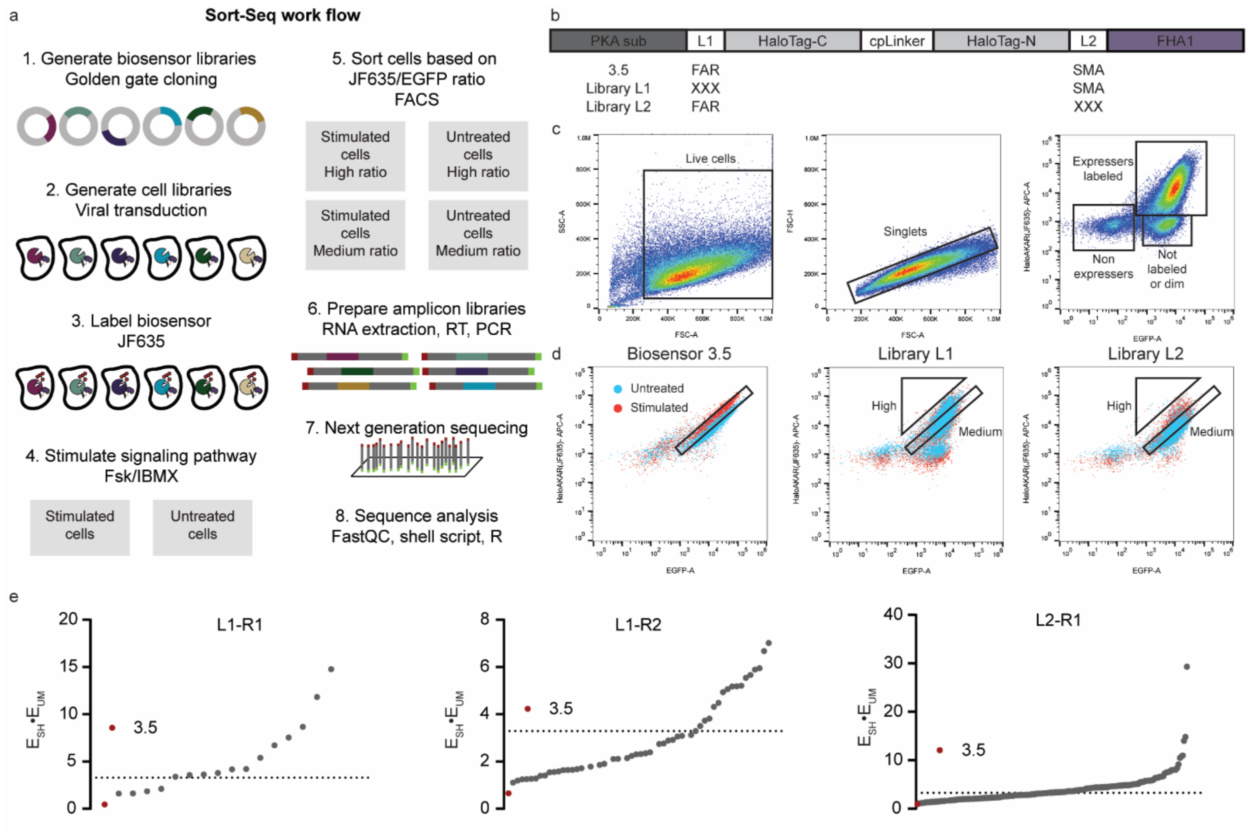

**Supplementary Figure S2**: Biosensor screening using Sort-Seq. **a**, Sort-Seq workflow. **b**, Domain structure of biosensor used in screening. Based on biosensor 3.5 (HaloAKAR1.0, Supplementary Table S1 and Supplementary Methods Table S1), we generated libraries L1 and L2 as indicated. **c**, Summary of FACS gating strategy. Live and singlet cells were gated as indicated. Within the live-singlet population, three subpopulations were visible: non-expressers; non-labeled/dim, which expresses EGFP but show no or little $JF_{635}$ (APC) fluorescence; or labeled expressers. Representative data are shown. **d**, Distribution of $JF_{635}$ (APC) and EGFP signals of the stimulated and untreated cells, along with the “High” and “Medium” gates used for sorting. Biosensor 3.5 (HaloAKAR1.0) was used as a reference to place the medium gate. Representative data are shown. **e**, Biosensor variants fulfilling the selection criteria ($E_{SH}>1$; $E_{UM}>1$; $E_{SM}<1$; $E_{UH}<1$) ranked according to the product of $E_{SH}$ and $E_{UM}$. Data for library L1 read 1 (R1) and read 2 (R2) as well as library L2 read 1 (R1) are given. Parental biosensor 3.5 (HaloAKAR1.0) is shown in red, and the cutoff used to select biosensors is given as a dotted line. (SH = Stimulated-high, UM = Untreated-medium, SM = Stimulated-medium, UH = Untreated-high).

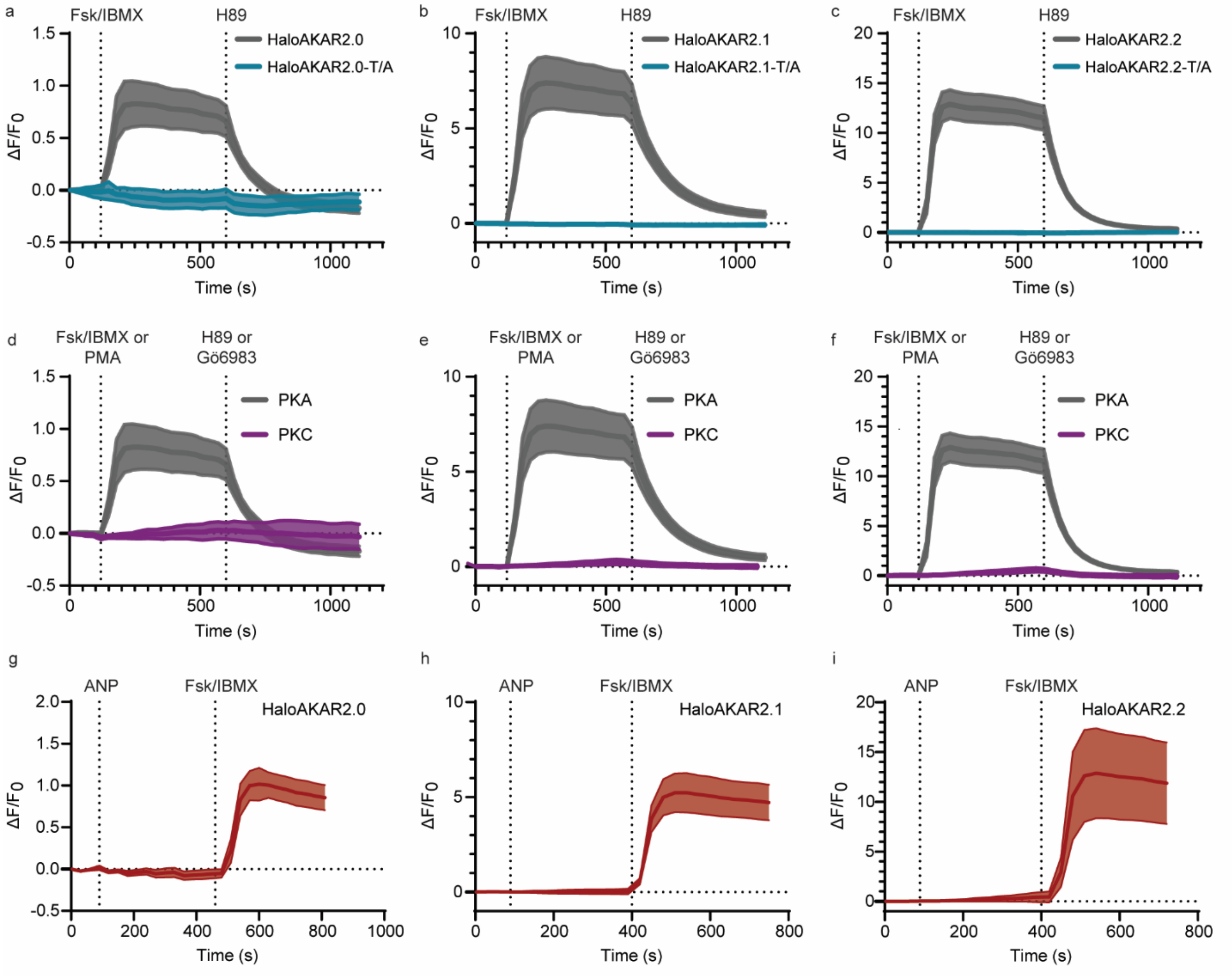

**Supplementary Figure S3**: Characterizing the best three biosensors for their specificity. **a-c**, Average time courses in HeLa cells expressing HaloAKAR2.0 (**a**), HaloAKAR2.1 (**b**), HaloAKAR2.2 (**c**) or the corresponding negative-control phospho-acceptor (T/A) mutants HaloAKAR2.0-T/A (**a**), HaloAKAR2.1-T/A (**b**), and HaloAKAR2.2-T/A (**c**) labeled with $JF_{635}$-CA and stimulated with 50 µM Fsk and 100 µM IBMX (Fsk/IBMX) and 20 µM H89 (n = 7, 10, 13, 6, 6, 5 cells 2.0, 2.1, 2.2, 2.0-T/A, 2.1-T/A, 2.2-T/A). **d-f**, Average time courses in HeLa cells expressing HaloAKAR2.0 (**d**), HaloAKAR2.1 (**e**), or HaloAKAR2.2 (**f**) labeled with $JF_{635}$-CA and stimulated with Fsk/IBMX and 20 µM H89 or 100 ng $mL^{-1}$ phorbol 12-myristate 13-acetate (PMA) and 1 µM Gö6983 (n = 7, 10, 13, 7, 8, 6 cells 2.0-PKA, 2.1-PKA, 2.2-PKA, 2.0-PKC, 2.1-PKC, 2.2-PKC) **g-i**, Average time courses in HeLa cells expressing HaloAKAR2.0 (**g**), HaloAKAR2.1 (**h**), or HaloAKAR2.2 (**i**) labeled with $JF_{635}$-CA and stimulated with 0.4 µM ANP and Fsk/IBMX (n = 7, 8, and 6 cells 2.0, 2.1, and 2.2). Time courses are representative of three replicates. Solid lines indicate mean response; shaded areas correspond to 95% confidence interval.

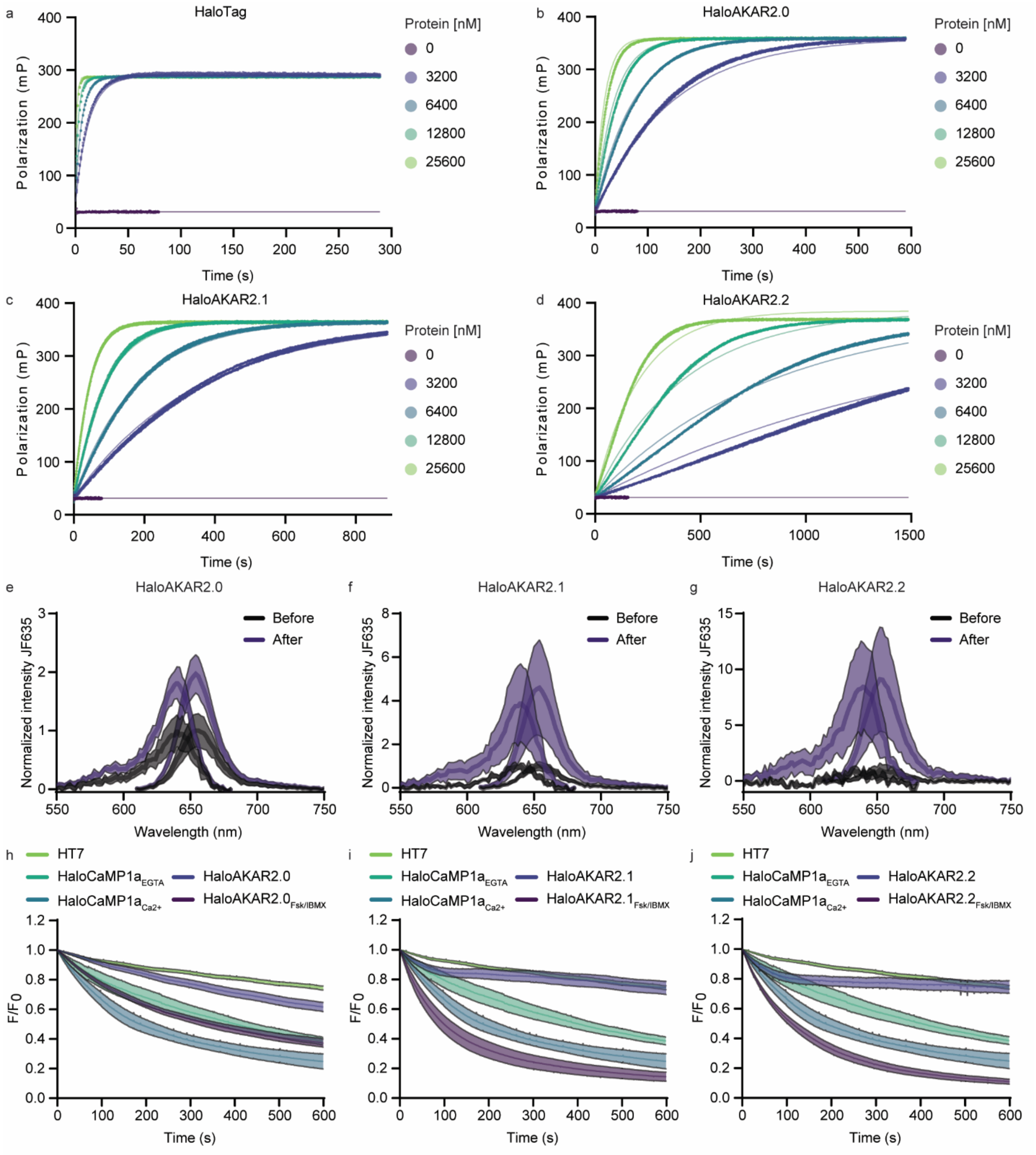

**Supplementary Figure S4**: *In vitro* and bleaching characterization of the best three biosensors. **a-d**, Labeling kinetics of HaloTag7 (**a**), HaloAKAR2.0 (**b**), HaloAKAR2.1 (**c**) or HaloAKAR2.2 (**d**) with Alexa488-CA (50 nM). Measured polarization traces (points) and fit curves (lines) are given for five different protein concentrations. For each concentration, three replicates were measured. Fit parameters and resulting rate constants can be found in Supplementary Table S6. **e-g**, Excitation and emission spectra of purified HaloAKAR2.0 (**e**), HaloAKAR2.1 (**f**) or HaloAKAR2.2 (**g**) labeled with $JF_{635}$-CA in the presence (purple) or absence (black) of $PKA_{cat}$. Solid lines indicate mean response; shaded areas correspond to standard deviation (n = 3 replicates). **h-j**, Bleaching curves of HaloAKAR2.0 (**h**), HaloAKAR2.1 (**i**), and HaloAKAR2.2

(**j**) labeled with $JF_{635}$-CA in the basal and stimulated (Fsk/IBMX) states. For comparison, bleaching of HaloTag7 and HaloCaMP1a in the calcium unbound (EGTA) and bound ($Ca^{2+}$) states, all labeled with $JF_{635}$-CA, are given. Survival fractions after bleaching for 600 s are given in Supplementary Table S8. Time courses are representative of three replicates. Solid lines indicate mean response; shaded areas correspond to standard deviation (n = 8, 9, 9, 11, 12, 9, 8, 9, and 9 cells HaloTag7, HaloCaMP1a-EGTA, HaloCaMP1a-$Ca^{2+}$, HaloAKAR2.0, HaloAKAR2.0-Fsk/IBMX, HaloAKAR2.1, HaloAKAR2.1-Fsk/IBMX, HaloAKAR2.2, and HaloAKAR2.2-Fsk/IBMX).

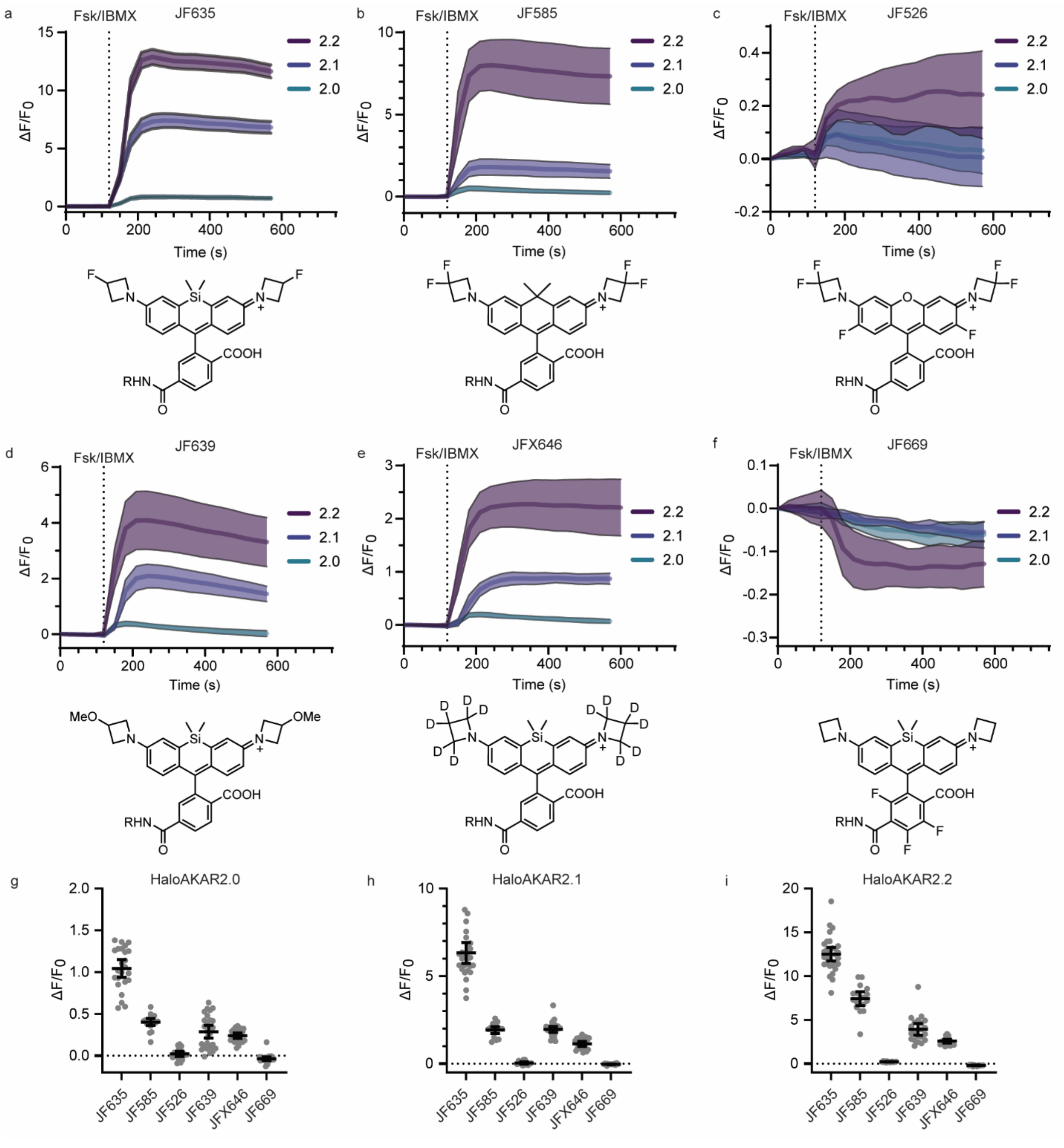

**Supplementary Figure S5**: HaloAKAR biosensor series labeled with rhodamine-based fluorophores of different spectral properties. **a-f,** Average time courses in HeLa cells expressing HaloAKAR2.0, HaloAKAR2.1, or HaloAKAR2.2 labeled with $JF_{635}$-CA (**a**), $JF_{585}$-CA (**b**), $JF_{526}$-CA (**c**), $JF_{639}$-CA (**d**) $JFX_{646}$-CA (**e**) or $JF_{669}$-CA (**f**) and stimulated with 50 µM Fsk and 100 µM IBMX (Fsk/IBMX). Time courses are representative of three replicates. Solid lines indicate mean response; shaded areas correspond to 95% confidence interval (a: n = 7, 10, 13; b: n = 6, 6, 6; c: n = 6, 6, 5; d: n = 9, 10, 6; e: n = 7, 5, 5; f: n = 8, 6, 4 cells 2.0, 2.1, 2.2). **g-i**, Maximum $\Delta F/F_0$ for HaloAKAR2.0 (**g**), HaloAKAR2.1 (**h**) and HaloAKAR2.2 (**i**) labeled with different rhodamine-based fluorophores. Individual data points are given along with mean and 95%

confidence interval (g: n = 24, 20, 20, 27, 24, 23 cells; h: n = 28, 17,13, 27, 22, 21 cells; i: n = 30, 17, 15, 21, 20, 15 cells from three replicates).

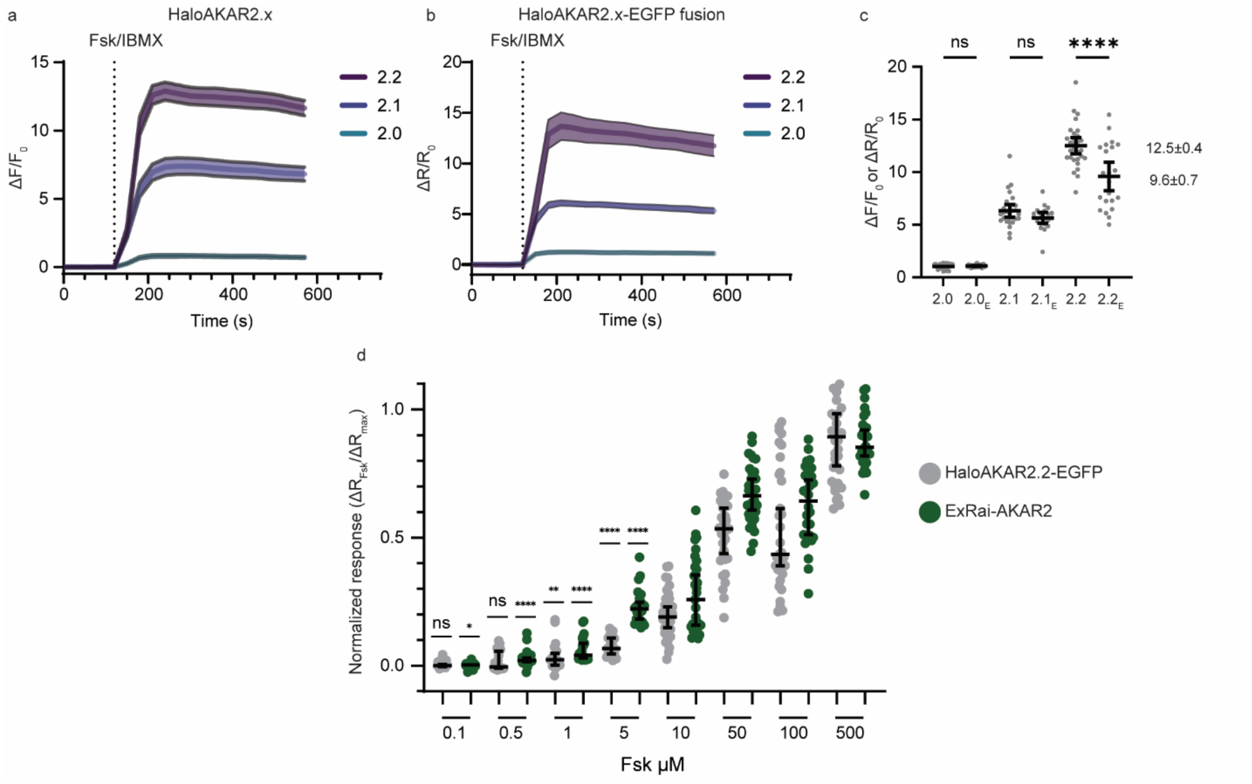

**Supplementary Figure S6**: Dose response of HaloAKAR2.2-EGFP and ExRaiAKAR2. **a-b,** Average time courses in HeLa cells expressing HaloAKAR2.0, HaloAKAR2.1, or HaloAKAR2.2 (**a**) or HaloAKAR2.0-EGFP, HaloAKAR2.1-EGFP, or HaloAKAR2.2-EGFP (**b**) labeled with $JF_{635}$-CA and stimulated with 50 µM Fsk and 100 µM IBMX (Fsk/IBMX). In (**b**) the $JF_{635}$/EGFP signal was used. Time courses are representative of three replicates. Solid lines indicate mean response; shaded areas correspond to 95% confidence interval (n = 7, 10, 13, 7, 9, 7 cells for 2.0, 2.1, 2.2, 2.0-EGFP, 2.1-EGFP, 2.2-EGFP, respectively). **c**, Maximum $\Delta F/F_0$ for HaloAKAR2.0, 2.1, 2.2 and the EGFP-fusions are given. Individual data points are shown along with mean and 95% confidence interval (n = 24, 28, 30, 20, 19, and 22 cells from three replicates, ***P < 0.0001, DFd = 141 one-way ANOVA followed by Holm Šidák's multiple-comparison test). **d**, Dose response measured in HeLa cells expressing either HaloAKAR2.2-EGFP labeled with $JF_{635}$-CA or ExRaiAKAR2. Cells were stimulated with the indicated doses of Fsk, followed by stimulation with 50 µM Fsk and 100 µM IBMX. HaloAKAR2.1-$JF_{635}$ showed a significant response to Fsk at concentrations as low as 1 µM. Currently, only ExRai-AKAR2, which is brighter in the basal state, can respond to lower doses (0.1 µM). Data are plotted as ($\Delta R$[Fsk])/($\Delta R_{max}$), where $\Delta R$[Fsk] is the difference in ratio recorded after addition of the indicated Fsk concentration, and $\Delta R_{max}$ is the difference in ratio after addition of Fsk and IBMX. Solid lines indicate the median and 95% confidence interval (n = 23, 26, 21, 23, 20, 21, 21, 24, 35, 31, 32, 34, 33, 30, 36, 28 cells from three replicates (left to right), **** P ≤ 0.0001, *** P ≤ 0.001, ** P ≤ 0.01, * P ≤ 0.05, ns P >0.05; vs. H0: median = 0.0; one-sample two-tailed Wilcoxon signed-rank test).

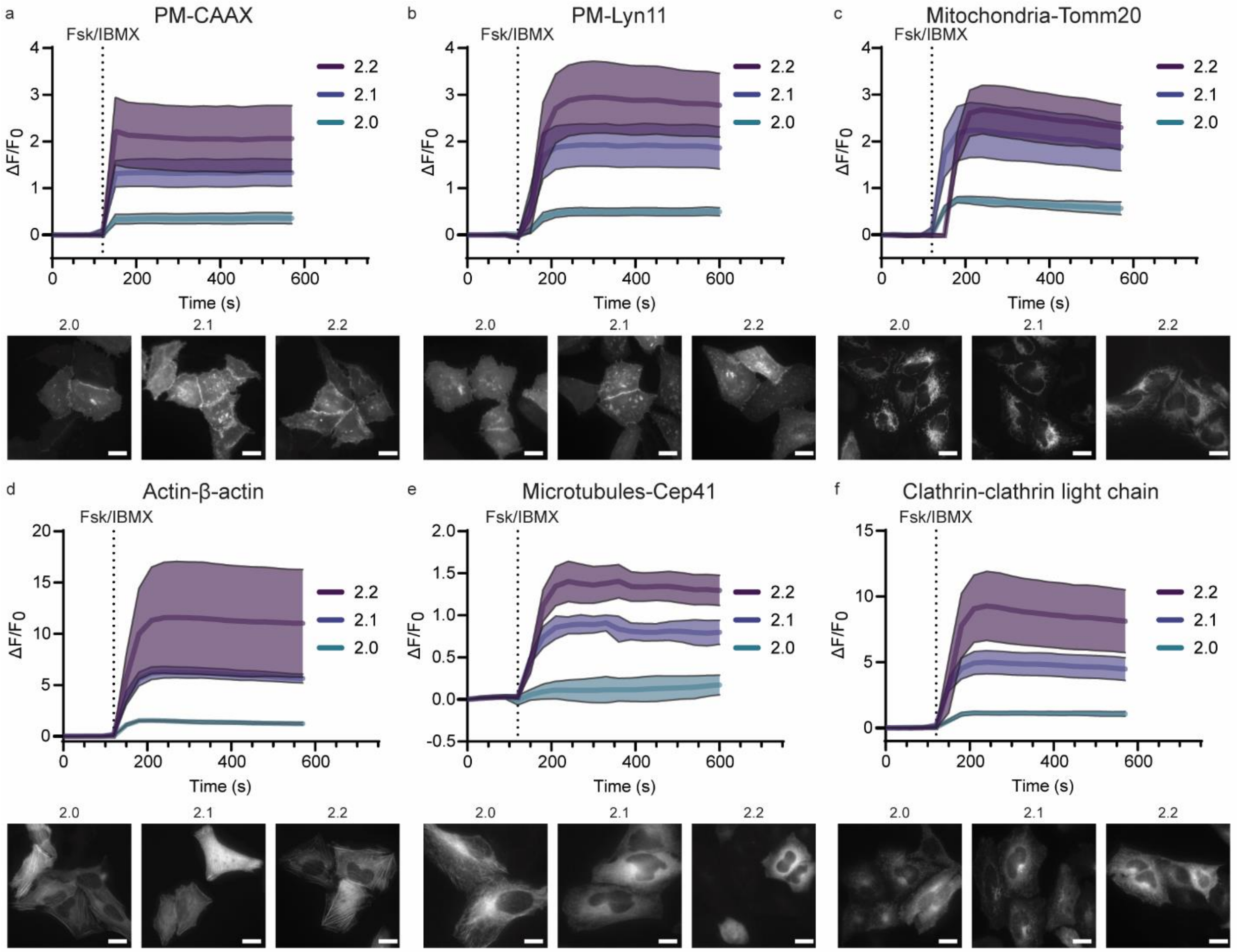

**Supplementary Figure S7**: Subcellularly targeted HaloAKAR biosensors. **a-b**, Plasma membrane targeting via fusion to the CAAX sequence of KRAS (**a**) or Lyn11 (**b**). **c**, Outer mitochondrial membrane targeting by fusing to Tomm20. **d**, Actin targeting via fusion with β-actin. **e**, Microtubule targeting via microtubule binding protein CEP41. **f**, Clathrin targeting via fusion with clathrin light chain. Average time courses of HeLa cells expressing targeted HaloAKAR2.0, HaloAKAR2.1, or HaloAKAR2.2 labeled with $JF_{635}$-CA and stimulated with 50 µM Fsk and 100 µM IBMX (Fsk/IBMX) are shown along with representative images after stimulation. Time courses and images are representative of three replicates. Solid lines indicate mean response; shaded areas correspond to 95% confidence interval (a: n = 8, 8, 7 cells; b: n = 7, 6, 4 cells; c: n = 7, 7, 5 cells; d: n = 5, 7, 4 cells; e: n = 6, 4, 5 cells; f: n = 5, 8, 6 cells). Scale bars, 20 µm.

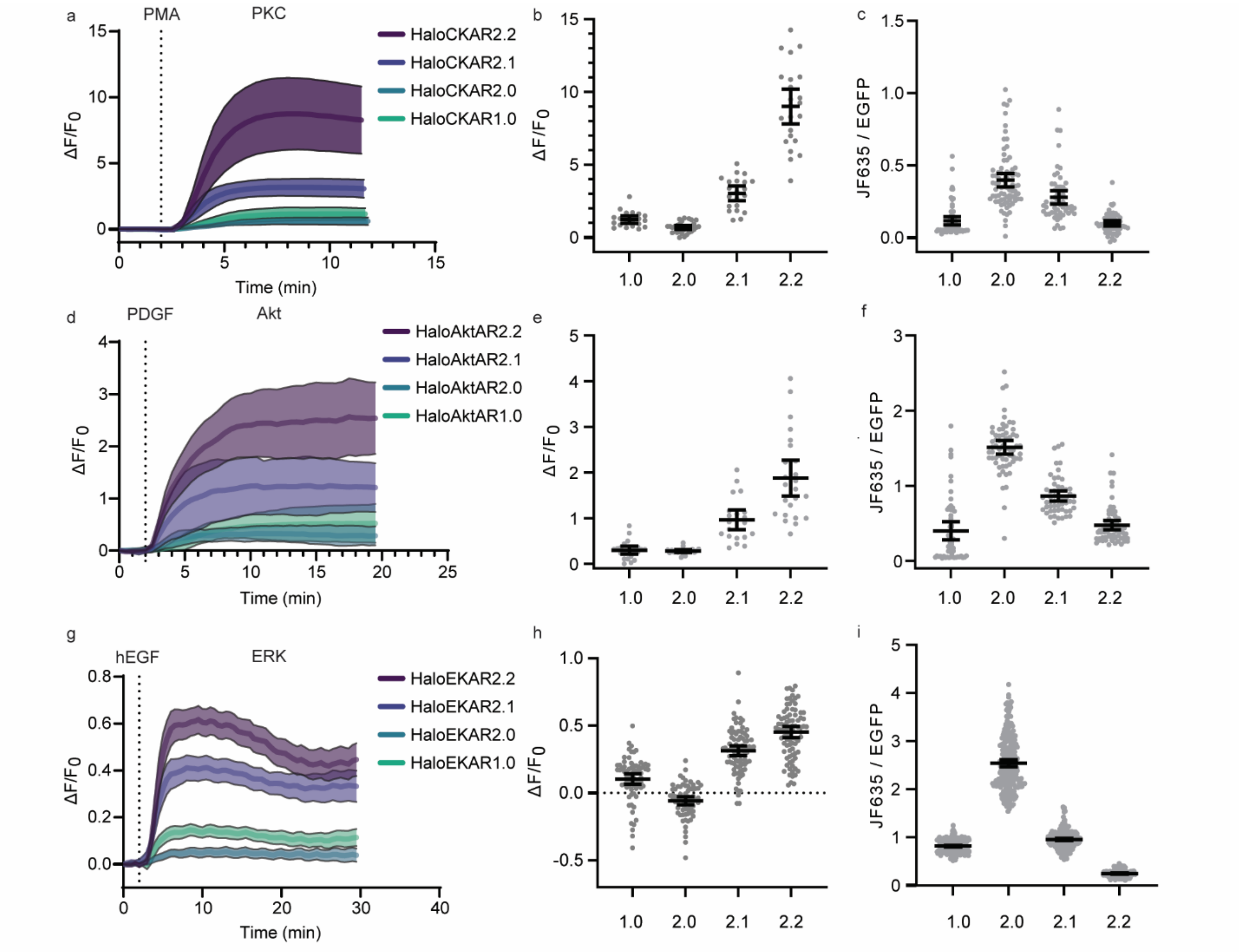

**Supplementary Figure S8**: The HaloAKAR design can be expanded to other kinases. **a-c**, Characterization of HeLa cells expressing HaloCKAR1.0, HaloCKAR2.0, HaloCKAR2.1, or HaloCKAR2.2 labeled with $JF_{635}$-CA and stimulated with 100 ng $mL^{-1}$ PMA. Average time courses (**a**) and quantification of dynamic range (**b**) and basal brightness (**c**) are given. **d-f**, Characterization of NIH3T3 cells expressing HaloAktAR1.0, HaloAktAR2.0, HaloAktAR2.1, or HaloAktAR2.2 labeled with $JF_{635}$-CA and stimulated with 50 µg $mL^{-1}$ PDGF. Average time courses (**d**), quantification of the maximum response after 8 min of stimulation (**e**), and the basal brightness (**f**) are given. **g-i**, Characterization of HEK293T cells expressing HaloEKAR1.0, HaloEKAR2.0, HaloEKAR2.1, or HaloEKAR2.2 labeled with $JF_{635}$-CA and stimulated with 100 ng $mL^{-1}$ hEGF. Average time courses (**g**), quantification of maximum response (**h**), and basal brightness (**i**) are given. Time courses are representative of three replicates. Solid lines indicate mean response; shaded areas correspond to 95% confidence interval (a-HaloCKAR: n = 7, 11, 6, 9; d-HaloAktKAR: n = 4, 4, 5, 8; g-HaloEKAR: n = 21, 24, 28, 35 cells 1.0, 2.0, 2.1, 2.2). For dynamic range and basal brightness, individual data points are given along with mean and 95% confidence interval. DR: b-HaloCKAR: 20, 30, 21, 23 cells from three replicates; e-HaloAktAR: 22, 21, 21, 24 cells from four replicates; h-HaloEKAR: 71, 68, 86, 88 cells from three replicates; Basal brightness: c-HaloCKAR: 61, 76, 55, 57 cells from nine fields of view (FOVs) and three replicates; f-HaloAktAR: 53, 61, 53, 64 cells from 12 FOVs and four replicates; i-HaloEKAR: 220, 232, 223, 168 cells from 9 FOVs and three replicates.

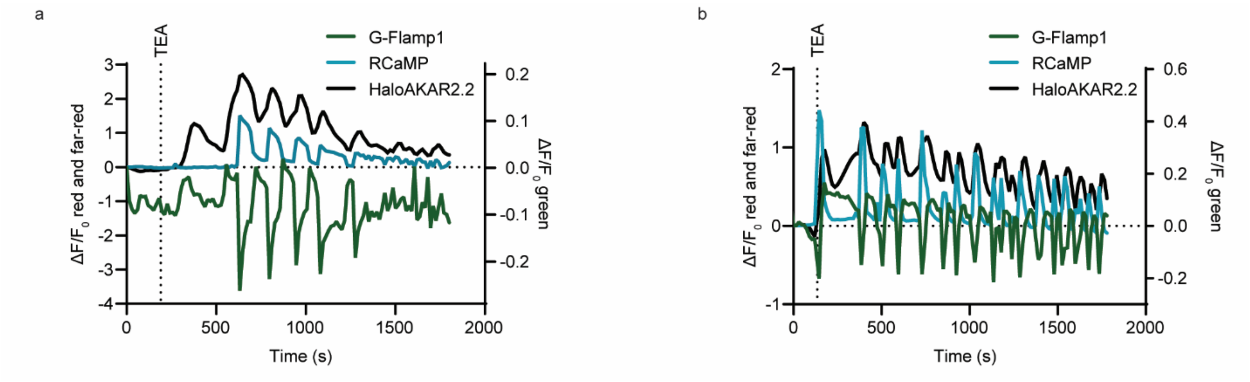

**Supplementary Figure S9**: Three-color multiplexed imaging of signaling cascades in MIN6 cells using single-fluorophore biosensors. **a-b**, Time course of G-Flamp1 (green), RCaMP (blue), and HaloAKAR2.2 (black) responses in single MIN6 cells stimulated with 20 mM tetraethylammonium chloride (TEA). Shown are representative traces are from three replicates.

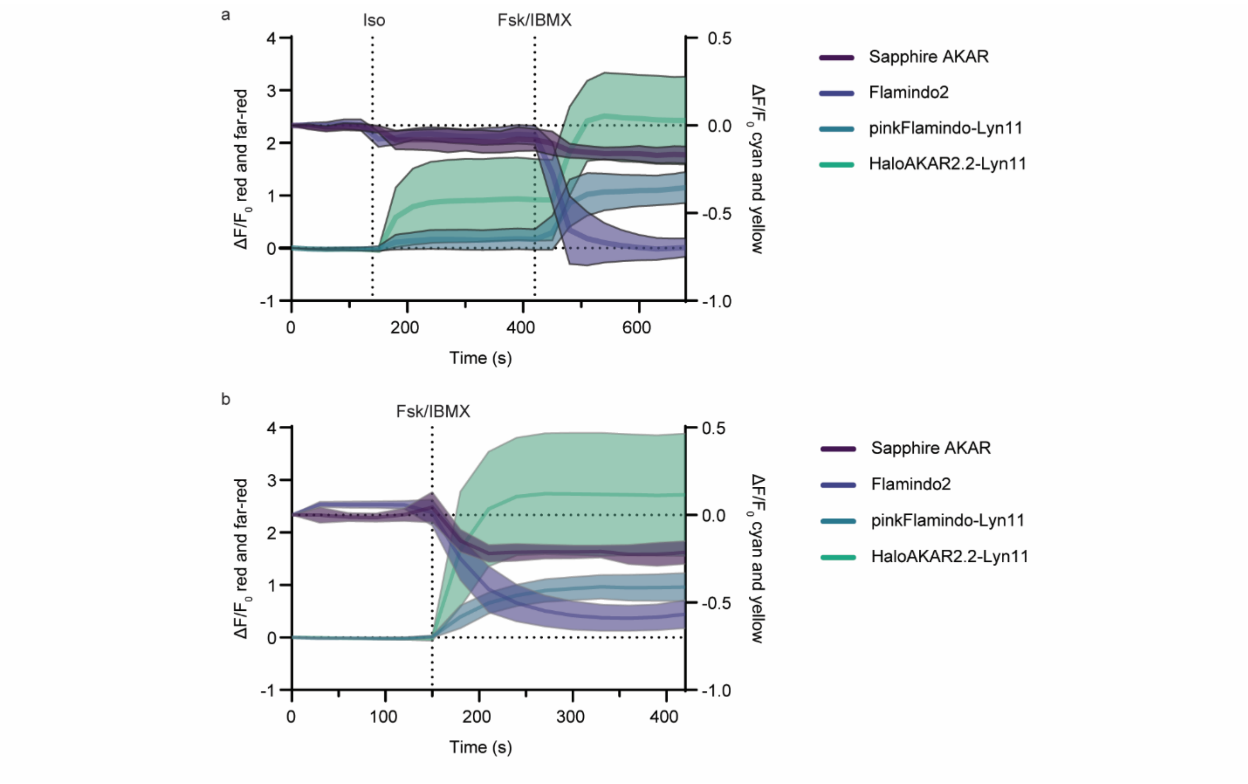

**Supplementary Figure S10**: Four-color multiplexed imaging of cell signaling in HeLa cells using single-fluorophore biosensors. **a-b**, Time courses of HeLa cells expressing negative-going sapphireAKAR (PKA activity), negative-going Flamindo2 (cAMP), pinkFlamindo-Lyn11 at the plasma membrane (cAMP), and HaloAKAR2.2-Lyn11-$JF_{635}$ at the plasma membrane (PKA activity) stimulated with (**a**) 1 µM isoproterenol (iso) followed by 50 µM Fsk and 100 µM IBMX (Fsk/IBMX) or (**b**) Fsk/IBMX directly. Time courses are representative of three replicates. Solid lines indicate mean response; shaded areas correspond to 95% confidence interval (n = 6 cells).

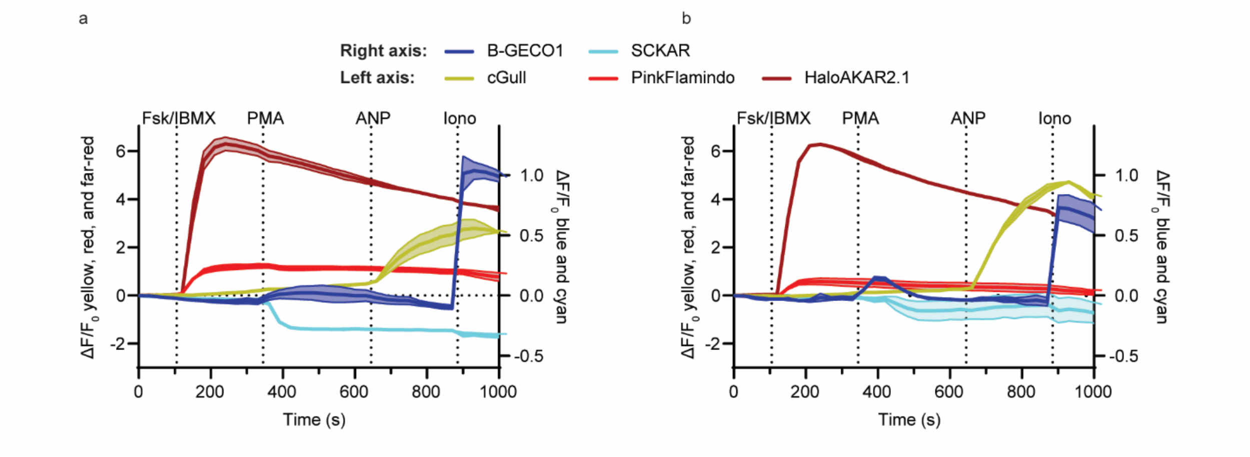

**Supplementary Figure S11**: Five-color multiplexed imaging of cell signaling in HeLa cells using single-fluorophore biosensors. **a-b**, Time courses of HeLa cells expressing B-GECO1 (blue, $Ca^{2+}$), negative-going sapphireCKAR (SCKAR, cyan, PKC activity), cGull (yellow, cGMP), pinkFlamindo (red, cAMP), and HaloAKAR2.1-$JF_{635}$ (dark-red, PKA activity) stimulated with 50 µM Fsk and 100 µM IBMX (Fsk/IBMX), 100 ng $mL^{-1}$ (PMA), 0.4 µM atrial natriuretic peptide (ANP), and 1 µM ionomycin (Iono). Time courses are representatives of seven replicates. Solid lines indicate mean response; shaded areas correspond to standard deviation (n = 2 cells).

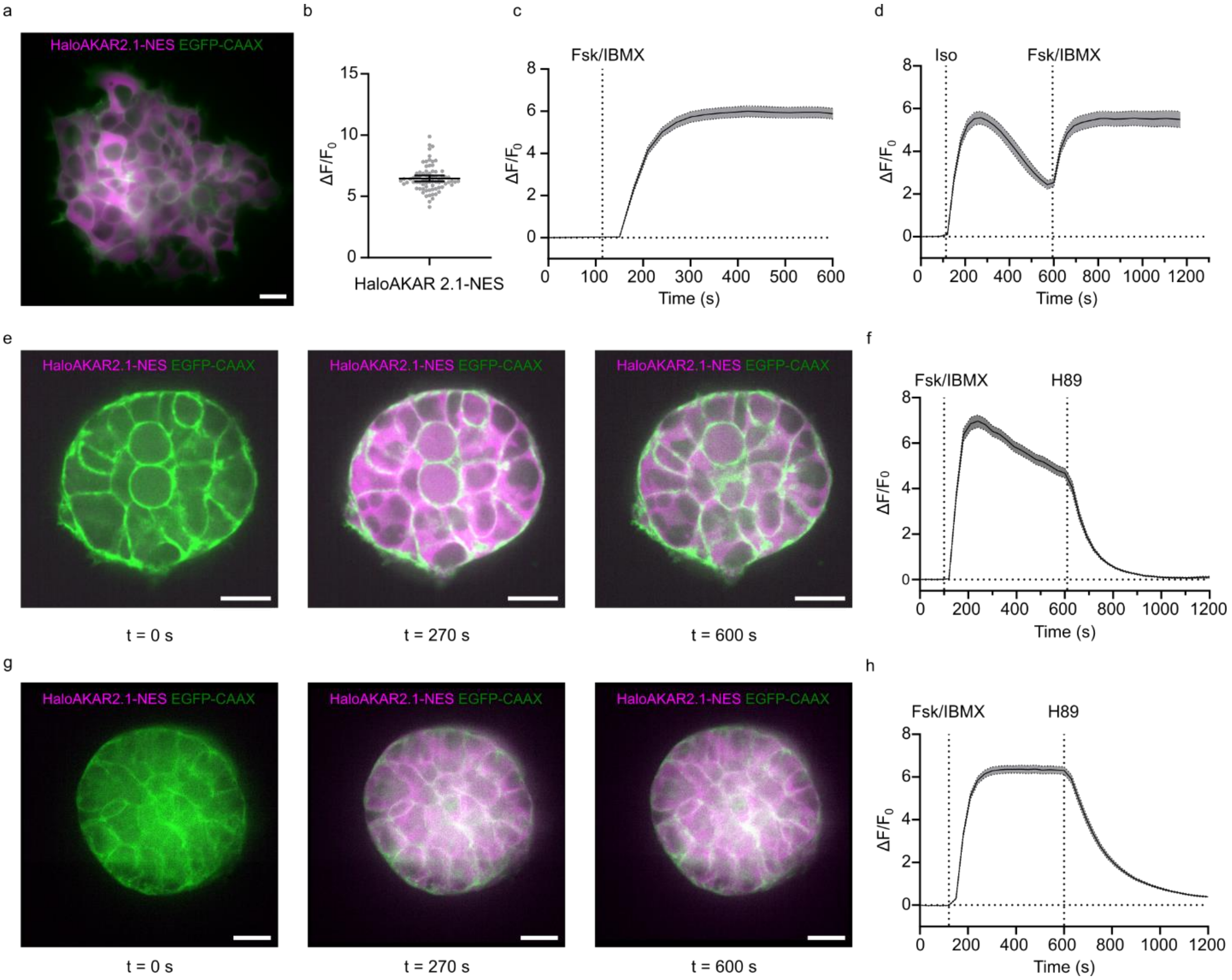

**Supplementary Figure S12**: Characterization of HaloAKAR2.1-NES in 2D and 3D culture. **a-d** Characterization of HEK293T cells stably expressing cytosolic HaloAKAR2.1 (HaloAKAR2.1-NES) and plasma membrane EGFP (EGFP-CAAX) labeled with $JF_{635}$-CA in 2D culture. A representative image overlay (EGFP: green, HaloAKAR: magenta) after Fsk/IBMX stimulation (**a**), the dynamic range ($\Delta F/F_0$) after Fsk/IBMX stimulation (**b**, n = 75 cells from three replicates), time courses after stimulation with Fsk/IBMX (**c**, n = 25 cells) or 1 µM isoproterenol (iso) followed by Fsk/IBMX (**d**, n = 28 cells) are given. **e-h**, Characterization of the same cell line in 3D spheroid culture. 4-d-old spheroids were labeled with $JF_{635}$-CA and imaged by spinning-disc confocal microscopy, acquiring images of the entire spheroid (**e-f**) or a single plane (**g-h**). Representative overlay images of a single plane (**e** and **g**) and the corresponding time course quantification (**f**: n = 27 cells, **h**: n = 20 cells) is given. Measuring z-stacks over the entire spheroid height led to some bleaching of the activated sensor, in contrast to the single-plane measurements (**g-h**). Time courses are representative of three replicates. Solid lines indicate mean response; shaded areas and error bars correspond to 95% confidence interval. Scale bars, 20 µm.

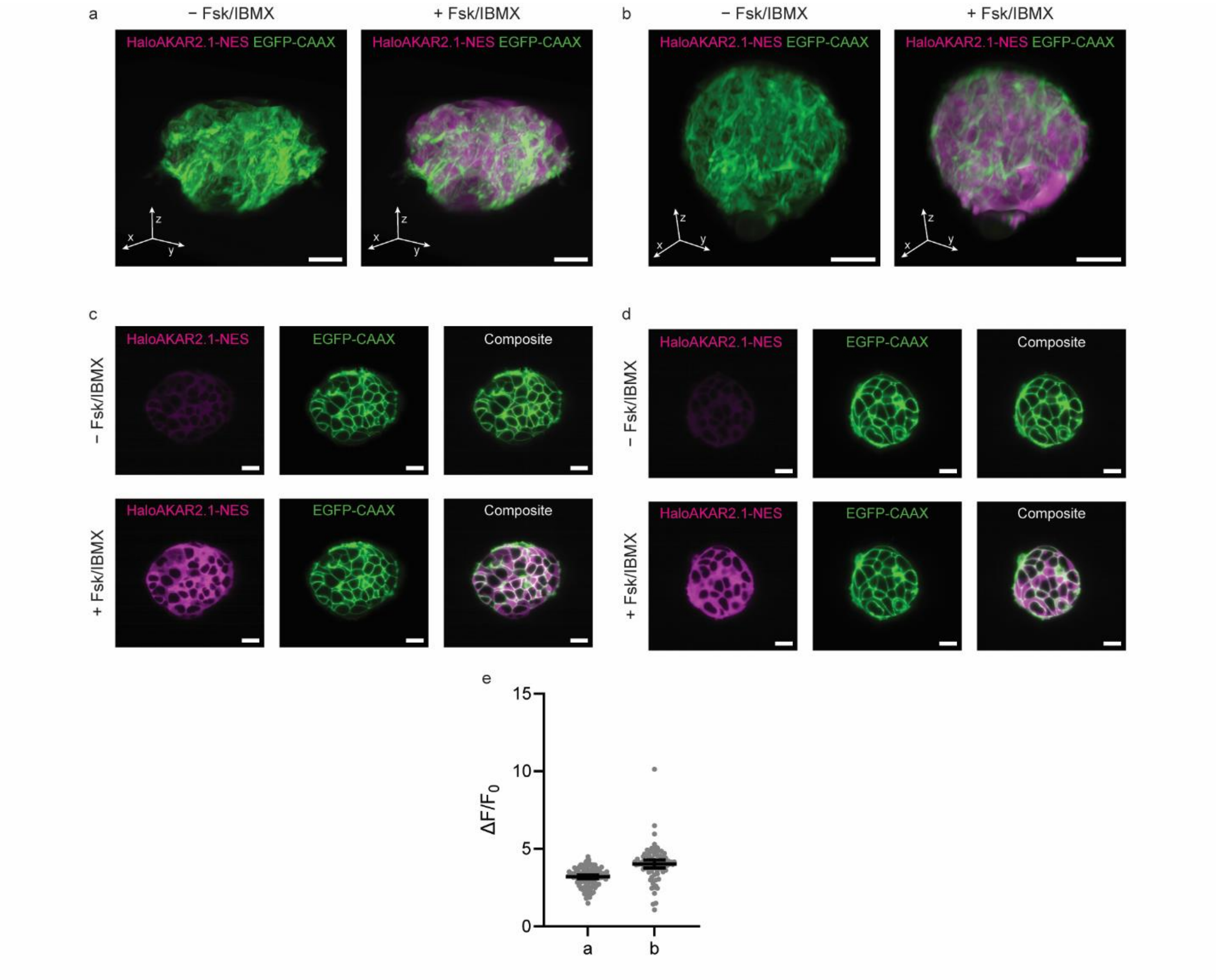

**Supplementary Figure S13**: Lattice-light-sheet imaging of HaloAKAR2.1-NES in HEK293T spheroids. **a-b** 3D renderings of 4-d-old spheroids generated from HEK293T cells stably expressing cytosolic HaloAKAR2.1 labeled with $JF_{635}$-CA (HaloAKAR2.1-NES-$JF_{635}$, magenta) and plasma membrane EGFP (EGFP-CAAX, green). Images of two replicates are given before and after Fsk/IBMX stimulation. **c-d**, Single plane of the spheroids before and after Fsk/IBMX stimulation clearly revealing individual cells. **e**, $\Delta F/F_0$ of individual cells in the spheroids before (**a**) and (**b**) after stimulation with Fsk/IBMX. Individual data points are given along with mean and 95% confidence interval (a: n = 108 cells, b: n = 78 cells). Scale bars, 20 µm.

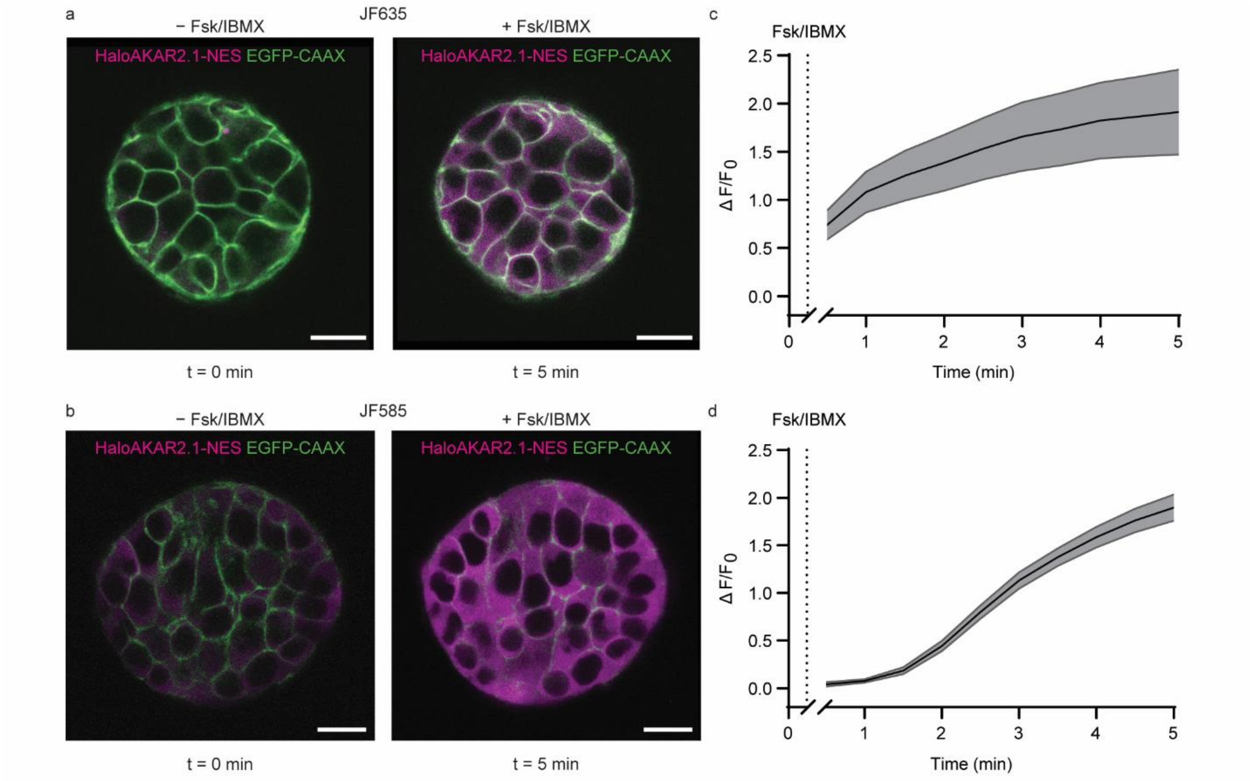

**Supplementary Figure S14**: Characterization of HaloAKAR2.1-NES in 3D culture by two-photon microscopy. **a-b,** Representative overlay images of 4-d-old spheroids generated from HEK293T cells stably expressing cytosolic HaloAKAR2.1 (HaloAKAR2.1-NES, magenta) and plasma membrane EGFP (EGFP-CAAX, green) labeled with $JF_{635}$-CA (**a**), or $JF_{585}$-CA (**b**). Images are shown before and after Fsk/IBMX stimulation. **c**-**d**, Corresponding time course quantification for spheroids are shown in (**c**, $JF_{635}$) and (**d**, $JF_{585}$). Time courses are representative of three replicates. Solid lines indicate mean response; shaded areas correspond to 95% confidence interval (c: n = 26 cells, d: n = 39 cells). Scale bars, 20 µm.

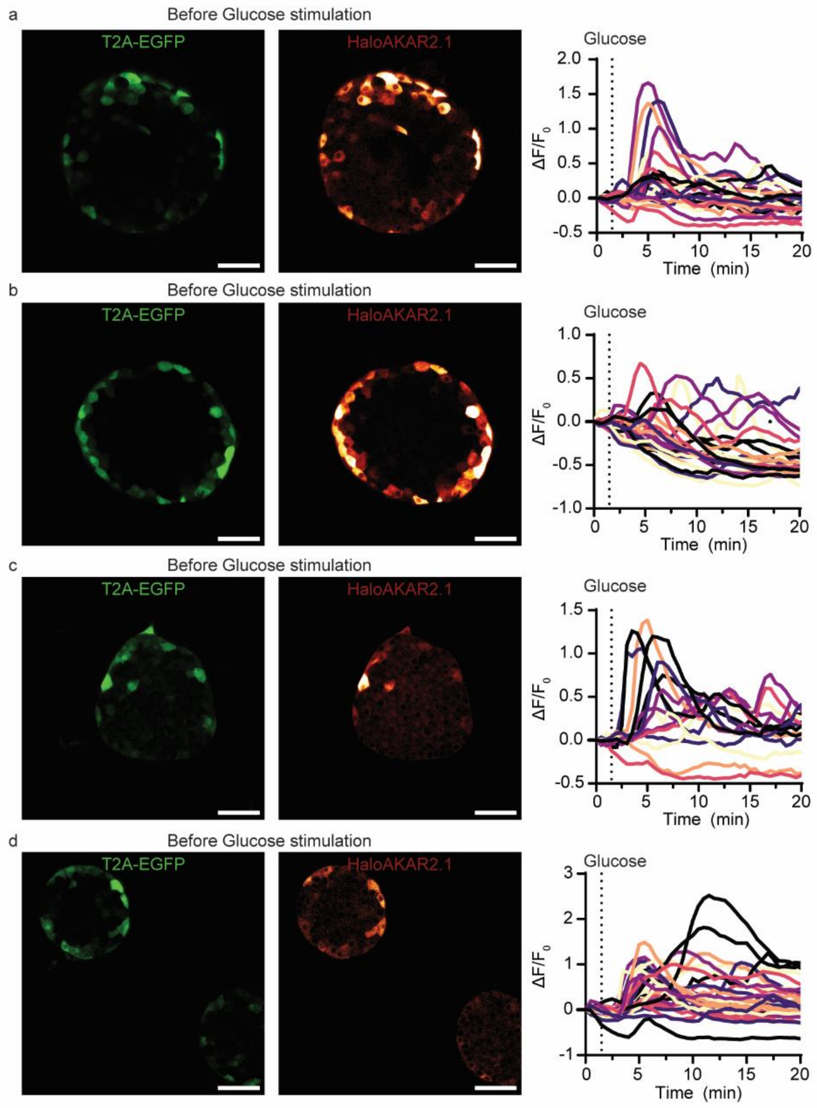

**Supplementary Figure S15**: Measurement of PKA activity in isolated mouse islets upon glucose stimulation using HaloAKAR2.1-$JF_{635}$. **a-d**, Images of EGFP (green) and $JF_{635}$-CA labeled HaloAKAR2.1 (red) are shown for time point t = 0 min along with time course of individual cells (n = 22, 25, 15, and 30 cells). Scale bars, 50 µm.

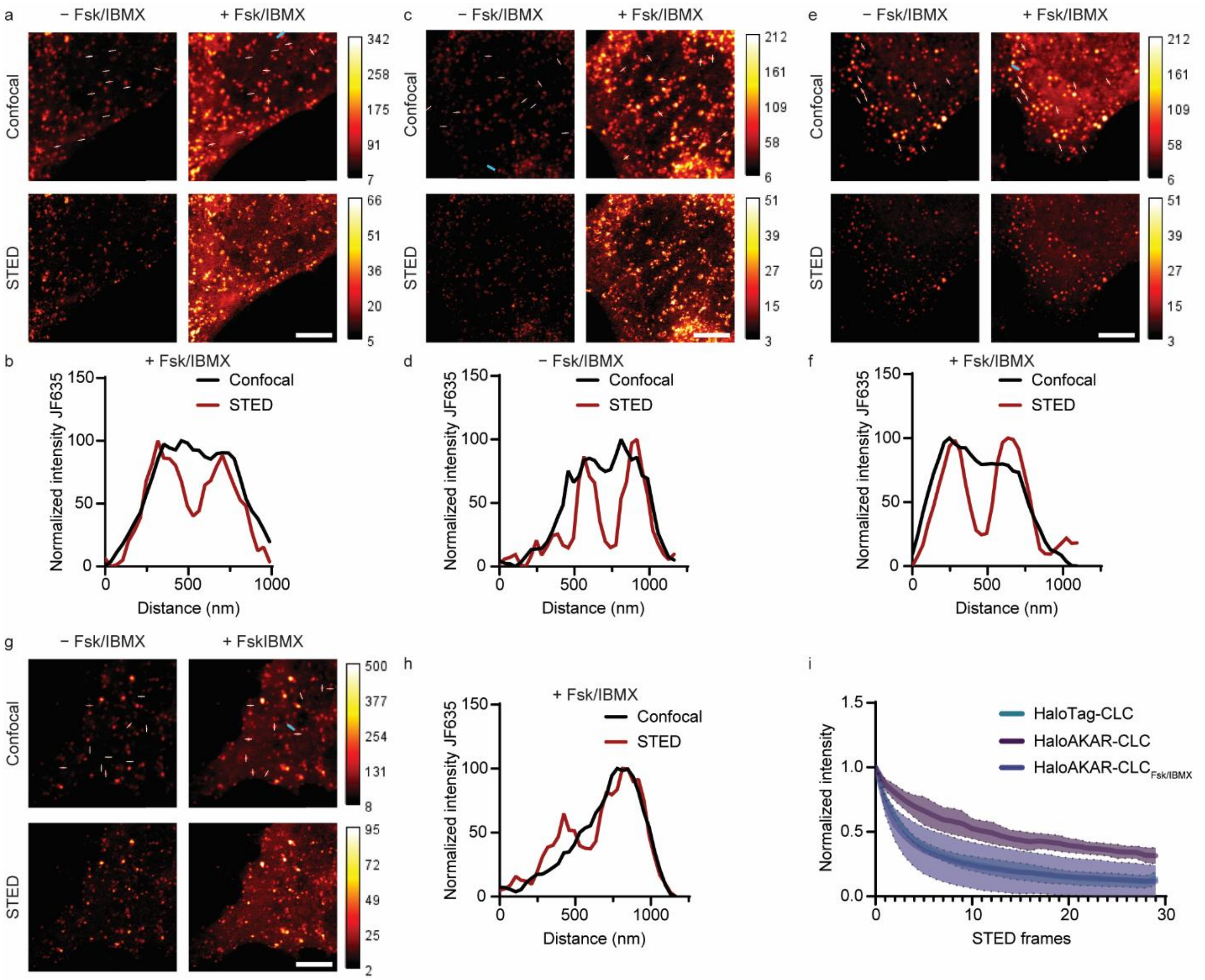

**Supplementary Figure S16**: STED microscopy measurements of HaloAKAR2.1-clathrin light chain (CLC) in live HeLa cells. **a-h**, Representative confocal and STED images of HeLa cells expressing HaloAKAR2.1-CLC labeled with $JF_{635}$-CA before and after Fsk/IBMX stimulation (**a**, **c**, **e**, **g**) along with representative line profiles across clathrin pits that can be resolved by STED but not confocal microscopy (**b**, **d**, **f**, **h**). Lines used to quantify the diameter of clathrin coated pits are indicated in the confocal images (white), as well as the lines used for the quantification in **b**, **d**, **f**, **h** (blue). **i**, Bleaching curves of HaloAKAR2.1-CLC labeled with $JF_{635}$-CA in the basal and stimulated (Fsk/IBMX) states (n = 3 cells from 3 replicates). For comparison, a bleaching curve for HaloTag7-CLC-$JF_{635}$ is given (n = 4 cells from 4 replicates). Solid lines indicate mean response; shaded areas correspond to standard deviation. Scale bars, 5 µm.

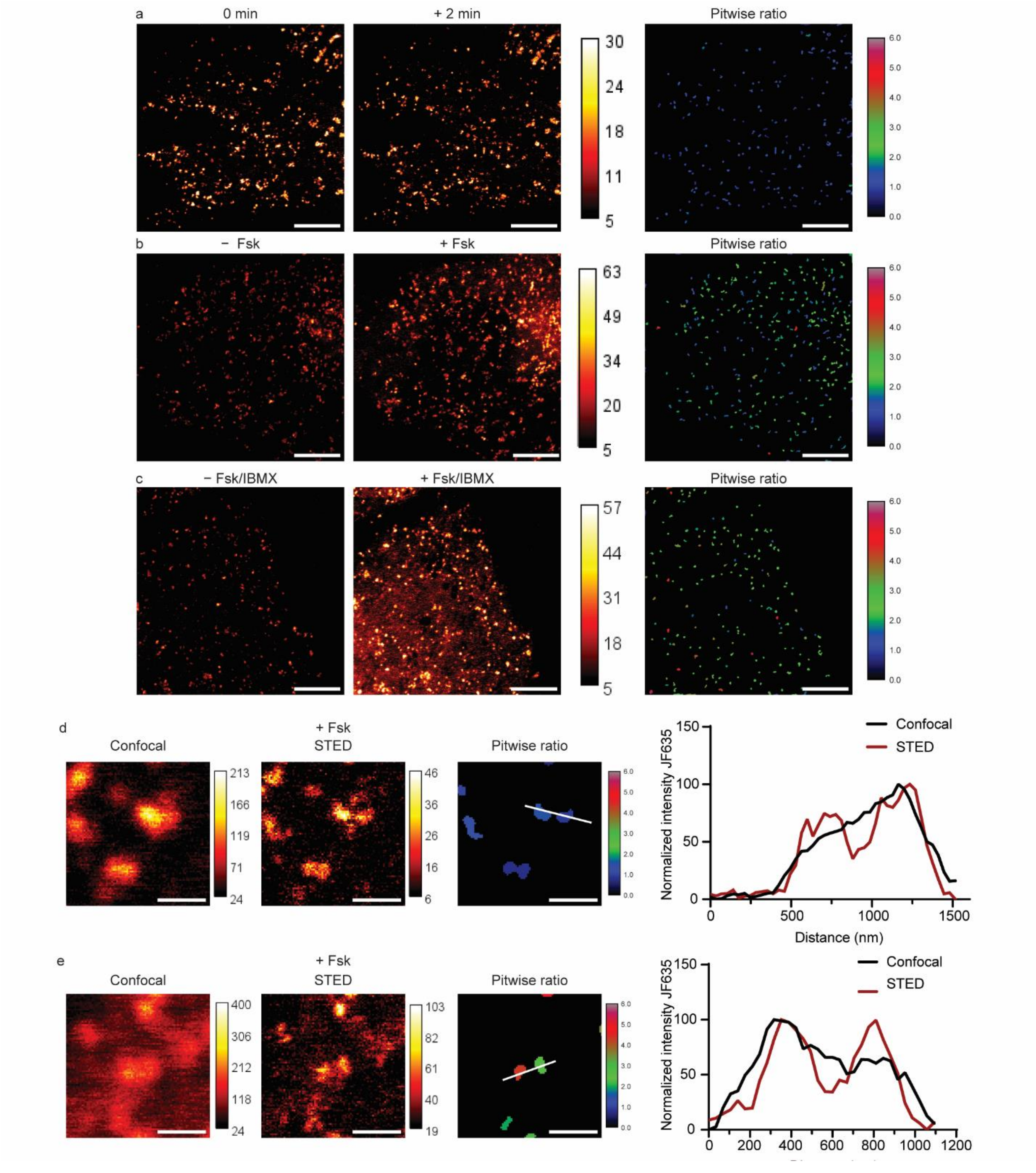

**Supplementary Figure S17**: STED microscopy measurements of HaloAKAR2.1-CLC in live HeLa cells. **a-c**, Representative STED images of HeLa cells expressing HaloAKAR2.1-CLC labeled with $JF_{635}$-CA before and after stimulation with no drug (**a**), 50 µM Fsk (**b**), 50 µM Fsk and 100 µM IBMX (**c**). Before and after images are shown along with pit-wise activity analysis showing the mean $F/F_0$ for pits present after drug treatment. Representative images for 5, 7 and 9 cells, respectively, are shown. Scale bars, 5 µm. **d**-**e**, Representative confocal, STED, and pitwise ratio images of HeLa cells expressing HaloAKAR2.1-CLC labeled with $JF_{635}$-CA after stimulation with 50 µM Fsk. Zoomed in images are shown, as well as the line profiles across the clathrin pits that can be resolved by STED but not confocal microscopy. Lines used for quantification are indicated in the ratio images. Scale bars, 500 nm.

### Supplementary Tables

**Supplementary Table S1**: Biosensor variants generated during four rounds of rational engineering. The responses ($\Delta F/F_0$) upon Fsk/IBMX stimulation, number of cells and FOV (corresponding to the number of replicates) are given. Variants in bold were characterized further. For later experiments Domain1 = Substrate and Domain2 = FHA1 were chosen.

| # | Domain1 | L1 | cpLink/Link | L2 | Domain2 | $\Delta F/F_0$±sem [%] | Cells | FOV |
|---|---|---|---|---|---|---|---|---|
| **1.1** | **Sub** | **AI** | **(GGTGGS)$_3$** | **SMH** | **FHA1** | **25.0±1.8** | **21** | **3** |
| **1.2** | **Sub** | **AI** | **EPTTGGS(GGTGGS)$_2$** | **SMH** | **FHA1** | **8.4±2.5** | **14** | **2** |
| **1.3** | Sub | FC | (GGTGGS)$_3$ | LLH | FHA1 | −30.1±2.1 | 15 | 2 |
| **1.4** | Sub | FC | EPTTGGS(GGTGGS)$_2$ | LLH | FHA1 | −6.8±1.5 | 14 | 2 |
| **1.5** | Sub | PEFAA | (GGTGGS)$_3$ | REFAR | FHA1 | −4.9±5.4 | 14 | 2 |
| **1.6** | Sub | PK | (GGTGGS)$_3$ | RFAR | FHA1 | 1.2±4.1 | 13 | 2 |
| **1.7** | **Sub** | **FAR** | **EPTTGGS(GGTGGS)$_2$** | **PEFAG** | **FHA1** | **19.3±1.4** | **11** | **2** |
| **1.8** | FAH1 | PEFAA | (GGTGGS)$_3$ | REFAR | Sub | −27.3±1.9 | 14 | 2 |
| **1.9** | FAH1 | PK | (GGTGGS)$_3$ | RFAR | Sub | −32.6±7.4 | 9 | 2 |
| **1.10** | FAH1 | FAR | EPTTGGS(GGTGGS)$_2$ | PEFAG | Sub | −11.1±2.1 | 12 | 2 |
| **1.11** | FAH1 | GT | (GGTGGS)$_3$ | KG | Sub | −17.2±2.8 | 11 | 2 |
| **1.12** | FAH1 | GT | (GGTGGS)$_3$ | SAGKPGSGEGSTKG | Sub | −20.6±2.1 | 7 | 1 |
| **1.13** | FAH1 | GT | EPTTGGS(GGTGGS)$_2$ | KG | Sub | −2.0±0.7 | 6 | 1 |
| **1.14** | FAH1 | GT | EPTTGGS(GGTGGS)$_2$ | SAGKPGSGEGSTKG | Sub | −21.0±0.8 | 6 | 1 |
| **1.15** | Halo150 | RMH | FHAI-14aa-sub | GGTGGSEL | Halo151 | −37.0±2.2 | 10 | 2 |
| **1.16** | Halo150 | - | FHAI-14aa-sub | GGT | Halo151 | −23.7±3.2 | 6 | 2 |
| **1.17** | Halo150 | RS | FHAI-EV-sub | GGSEL | Halo151 | −50.7±6.1 | 3 | 1 |
| **2.1** | Sub | FAR | (GGTGGS)$_3$ | PEFAG | FHA1 | 9.6±0.6 | 8 | 1 |
| **2.2** | **Sub** | **FAR** | **(GGTGGS)$_3$** | **SMH** | **FHA1** | **81.8±4.0** | **17** | **2** |
| **2.3** | **Sub** | **FAR** | **EPTTGGS(GGTGGS)$_2$** | **SMH** | **FHA1** | **76.4±3.3** | **13** | **2** |
| **2.4** | Sub | AI | (GGTGGS)$_3$ | PEFAG | FHA1 | −2.4±1.3 | 10 | 1 |
| **2.5** | Sub | AI | EPTTGGS(GGTGGS)$_2$ | PEFAG | FHA1 | 5.8±2.4 | 7 | 1 |
| **2.6** | Sub | AA | (GGTGGS)$_3$ | SMH | FHA1 | −17.3±0.9 | 8 | 1 |
| **2.7** | Sub | AI | (GGTGGS)$_3$ | AMH | FHA1 | −30.1±1.0 | 6 | 1 |
| **2.8** | Sub | AI | (GGTGGS)$_3$ | SAH | FHA1 | −10.4±1.3 | 6 | 1 |
| **2.9** | Sub | AI | (GGTGGS)$_3$ | SMA | FHA1 | −25.8±1.0 | 6 | 1 |
| **2.10** | Sub | AAR | EPTTGGS(GGTGGS)$_2$ | PEFAG | FHA1 | −15.8±1.3 | 10 | 1 |
| **2.11** | **Sub** | **FAA** | **EPTTGGS(GGTGGS)$_2$** | **PEFAG** | **FHA1** | **60.0±2.8** | **18** | **2** |
| **2.12** | **Sub** | **WI** | **(GGTGGS)$_3$** | **SMH** | **FHA1** | **25.5±1.1** | **17** | **2** |
| **2.13** | Sub | WAR | EPTTGGS(GGTGGS)$_2$ | PEFAG | FHA1 | 5.2±2.7 | 8 | 1 |
| **2.14** | **Sub** | **FWR** | **EPTTGGS(GGTGGS)$_2$** | **PEFAG** | **FHA1** | **33.2±2.2** | **17** | **2** |
| **2.15** | Sub | FAR | EPTTGGS(GGTGGS)$_2$ | PEWAG | FHA1 | 21.4±1.0 | 7 | 1 |
| **3.1** | Sub | AAR | EPTTGGS(GGTGGS)$_2$ | SMH | FHA1 | 30.1±2.0 | 8 | 1 |
| **3.2** | Sub | FAA | EPTTGGS(GGTGGS)$_2$ | SMH | FHA1 | 37.2±3.4 | 8 | 1 |
| **3.3** | Sub | FAR | EPTTGGS(GGTGGS)$_2$ | AMH | FHA1 | 49.9±1.8 | 8 | 1 |
| **3.4** | Sub | FAR | EPTTGGS(GGTGGS)$_2$ | SAH | FHA1 | 51.5±2.0 | 8 | 1 |
| **3.5** | **Sub** | **FAR** | **EPTTGGS(GGTGGS)$_2$** | **SMA** | **FHA1** | **146.9±4.5** | **14** | **2** |
| **3.6** | **Sub** | **FWR** | **EPTTGGS(GGTGGS)$_2$** | **SMH** | **FHA1** | **84.2±4.0** | **14** | **2** |
| **3.7** | **Sub** | **AFAR** | **EPTTGGS(GGTGGS)$_2$** | **SMH** | **FHA1** | **112.2±4.2** | **16** | **2** |

| | | | | | | | | |
|---|---|---|---|---|---|---|---|---|
| **3.8** | **Sub** | **FARA** | **EPTTGGS(GGTGGS)$_2$** | **SMH** | **FHA1** | **71.4±3.1** | **14** | **2** |
| **3.9** | Sub | FAR | EPTTGGS(GGTGGS)$_2$ | SM | FHA1 | 33.3±1.7 | 8 | 1 |
| **3.10** | **Sub** | **FAR** | **EPTTGGS(GGTGGS)$_2$** | **SMHA** | **FHA1** | **69.3±1.8** | **14** | **2** |
| **3.11** | Sub | AR | EPTTGGS(GGTGGS)$_2$ | SMH | FHA1 | 17.6±3.1 | 7 | 1 |
| **3.12** | Sub | FA | EPTTGGS(GGTGGS)$_2$ | SMH | FHA1 | 42.7±2.0 | 6 | 1 |
| **3.13** | **Sub** | **FAR** | **EPTTGGS(GGTGGS)$_2$** | **MH** | **FHA1** | **56.8±2.9** | **14** | **2** |
| **3.14** | Sub | FAR | EPTTGGS(GGTGGS)$_2$ | SM | FHA1 | 44.9±3.2 | 7 | 1 |
| **3.15** | Sub | FAR −1 | EPTTGGS(GGTGGS)$_2$ | SMH | FHA1 | 8.9±1.1 | 6 | 1 |
| **3.16** | Sub | FAR | EPTTGGS(GGTGGS)$_2$ | −1 SMH | FHA1 | −42.5±1.9 | 7 | 1 |
| **3.17** | Sub | FAR −2 | EPTTGGS(GGTGGS)$_2$ | SMH | FHA1 | −62.0±2.6 | 7 | 1 |
| **3.18** | Sub | FAR | EPTTGGS(GGTGGS)$_2$ | −2 SMH | FHA1 | 25.8±4.5 | 7 | 1 |
| **3.19** | Sub | FR | EPTTGGS(GGTGGS)$_2$ | SMH | FHA1 | 33.6±2.6 | 6 | 1 |
| **3.20** | Sub | FAR | EPTTGGS(GGTGGS)$_2$ | SH | FHA1 | 54.1±1.4 | 7 | 1 |
| **3.21** | **Sub** | **FAR** | **EPTTGGS(GGTGGS)$_2$** | **MR** | **FHA1** | **60.8±3.7** | **15** | **2** |
| **4.1** | Sub | AAR | EPTTGGS(GGTGGS)$_2$ | SMA | FHA1 | 36.2±3.6 | 9 | 1 |
| **4.2** | Sub | FAA | EPTTGGS(GGTGGS)$_2$ | SMA | FHA1 | 36.2±4.8 | 7 | 1 |
| **4.3** | Sub | FAR | EPTTGGS(GGTGGS)$_2$ | AMA | FHA1 | 56.2±2.1 | 9 | 1 |
| **4.4** | Sub | FAR | EPTTGGS(GGTGGS)$_2$ | SAA | FHA1 | 62.5±3.8 | 8 | 1 |
| **4.5** | Sub | FWR | EPTTGGS(GGTGGS)$_2$ | SMA | FHA1 | 66.8±3.8 | 8 | 1 |
| **4.6** | **Sub** | **AFAR** | **EPTTGGS(GGTGGS)$_2$** | **SMA** | **FHA1** | **155.4±7.9** | **16** | **2** |
| **4.7** | Sub | FARA | EPTTGGS(GGTGGS)$_2$ | SMA | FHA1 | 43.8±1.5 | 10 | 1 |
| **4.8** | Sub | FAR | EPTTGGS(GGTGGS)$_2$ | SMAA | FHA1 | 58.0±2.2 | 8 | 1 |
| **4.9** | Sub | FAR | EPTTGGS(GGTGGS)$_2$ | MA | FHA1 | 32.5±2.4 | 9 | 1 |
| **4.10** | Sub | FAR | EPTTGGS(GGTGGS)$_2$ | SA | FHA1 | 41.6±3.7 | 9 | 1 |
| **4.11** | Sub | FAR | EPTTGGS(GGTGGS)$_2$ | SMA | FHA1 | 40.2±1.8 | 10 | 1 |
| **4.12** | Sub | FAR | EPTTGGS(GGTGGS)$_2$ | SMA | FHA1 | 57.9±5.7 | 8 | 1 |
| **4.13** | Sub | FAR | EPTTGGS(GGTGGS)$_2$ | SMA | FHA1 | 52.1±4.4 | 9 | 1 |
| **4.14** | Sub | FAR | EPTTGGS(GGTGGS)$_2$ | SMA | −1 FHA1 | 29.0±3.0 | 7 | 1 |
| **4.15** | Sub | FAR | EPTTGGS(GGTGGS)$_2$ | SMA | −2 FHA1 | 38.2±2.8 | 6 | 1 |
| **4.16** | Sub | FAR | EPTTGGS(GGTGGS)$_2$ | SMA | −3 FHA1 | 29.3±5.1 | 6 | 1 |
| **4.17** | Sub | PEFAA | EPTTGGS(GGTGGS)$_2$ | SMA | FHA1 | 12.0±1.4 | 10 | 1 |
| **4.18** | Sub | FAR | EPTTGGS(GGTGGS)$_2$ | REFAR | FHA1 | 7.6±1.8 | 8 | 1 |

**Supplementary Table S2**: Best biosensor variants from each round of rational engineering. The response ($\Delta F/F_0$) upon Fsk/IBMX stimulation and the basal brightness ($JF_{635}$/EGFP) are given. Additionally, number of cells and FOV (corresponding to the number of replicates) are given. The two biosensors with the highest $\Delta F/F_0$ are highlighted in bold/italic and the four biosensors with the highest basal brightness in bold. For all biosensors, Domain 1 = Substrate and Domain 2 = FHA1. For subsequent experiments, cpLink = EPTTGGS(GGTGGS)$_2$ was chosen. The lengths of linkers L1 and L2 were fixed to three amino acids each.

| # | L1 | cpLink | L2 | $\Delta F/F_0$±sem [%] | Cells | FOV | $JF_{635}$/EGFP±sem | Cells | FOV |
|---|---|---|---|---|---|---|---|---|---|
| **1.1** | **AI** | **(GGTGGS)$_3$** | **SMH** | **25.0±1.8** | **21** | **3** | **0.375±0.017** | **27** | **3** |
| **1.2** | **AI** | **EPTTGGS(GGTGGS)$_2$** | **SMH** | **8.4±2.5** | **14** | **2** | **0.793±0.040** | **24** | **3** |
| 1.7 | FAR | EPTTGGS(GGTGGS)$_2$ | PEFAG | 19.3±1.4 | 11 | 2 | 0.248±0.009 | 26 | 3 |
| 2.2 | FAR | (GGTGGS)$_3$ | SMH | 81.8±4.0 | 17 | 2 | 0.182±0.015 | 25 | 3 |
| 2.3 | FAR | EPTTGGS(GGTGGS)$_2$ | SMH | 76.4±3.3 | 13 | 2 | 0.112±0.008 | 26 | 3 |
| 2.11 | FAA | EPTTGGS(GGTGGS)$_2$ | PEFAG | 60.0±2.8 | 18 | 2 | 0.226±0.10 | 29 | 3 |
| **2.12** | **WI** | **(GGTGGS)$_3$** | **SMH** | **25.5±1.1** | **17** | **2** | **1.105±0.047** | **24** | **3** |
| 2.14 | FWR | EPTTGGS(GGTGGS)$_2$ | PEFAG | 33.2±2.2 | 17 | 2 | 0.127±0.018 | 18 | 3 |
| ***3.5*** | ***FAR*** | ***EPTTGGS(GGTGGS)$_2$*** | ***SMA*** | ***146.9±4.5*** | ***14*** | ***2*** | ***0.108±0.003*** | ***32*** | ***3*** |
| 3.6 | FWR | EPTTGGS(GGTGGS)$_2$ | SMH | 84.2±4.0 | 14 | 2 | 0.065±0.006 | 24 | 3 |
| 3.7 | AFAR | EPTTGGS(GGTGGS)$_2$ | SMH | 112.2±4.2 | 16 | 2 | 0.177±0.014 | 28 | 3 |
| **3.8** | **FARA** | **EPTTGGS(GGTGGS)$_2$** | **SMH** | **71.4±3.1** | **14** | **2** | **0.300±0.012** | **25** | **3** |
| 3.10 | FAR | EPTTGGS(GGTGGS)$_2$ | SMHA | 69.3±1.8 | 14 | 2 | 0.098±0.007 | 24 | 3 |
| 3.13 | FAR | EPTTGGS(GGTGGS)$_2$ | MH | 56.8±2.9 | 14 | 2 | 0.113±0.006 | 27 | 3 |
| 3.21 | FAR | EPTTGGS(GGTGGS)$_2$ | MR | 60.8±3.7 | 15 | 2 | 0.126±0.004 | 26 | 3 |
| ***4.6*** | ***AFAR*** | ***EPTTGGS(GGTGGS)$_2$*** | ***SMA*** | ***155.4±7.9*** | ***16*** | ***2*** | ***0.126±0.004*** | ***32*** | ***3*** |

**Supplementary Table S3**: Biosensor variants selected from sequencing data and validated by fluorescence microscopy. The response ($\Delta F/F_0$) upon Fsk/IBMX stimulation and the basal brightness ($JF_{635}$/EGFP) are given. Additionally, number of cells and FOV are given. For $\Delta F/F_0$, FOV correspond to replicates; for basal brightness, six FOVs were measured from two replicates. The parental biosensor 3.5 = HaloAKAR1.0 is given in the first row. The biosensors selected for combination with each other are highlighted in bold. Biosensors were selected as follows: high dynamic range but low basal brightness, medium dynamic range and medium basal brightness, or low dynamic range and high basal brightness. Five biosensors were selected from library L1 and three from library L2.

| # | L1 | L2 | $\Delta F/F_0$±sem [%] | Cells | FOV | $JF_{635}$/EGFP±sem | Cells | FOV |
|---|---|---|---|---|---|---|---|---|
| **3.5** | **FAR** | **SMA** | 129.8±9.9 | 29 | **6** | 0.231±0.014 | 73 | 18 |
| **5.1** | **QMR** | **SMA** | **67.0±2.8** | **13** | **2** | **1.096±0.032** | **36** | **6** |
| **5.2** | **CKK** | **SMA** | **101.3±4.8** | **15** | **2** | **0.784±0.047** | **37** | **6** |
| **5.3** | **MKR** | **SMA** | **301.6±22.0** | **10** | **2** | **0.436±0.062** | **29** | **6** |
| 5.4 | YLH | SMA | 214.5±15.6 | 13 | 2 | 0.299±0.010 | 30 | 6 |
| **5.5** | **MRR** | **SMA** | **554.0±23.1** | **12** | **2** | **0.345±0.009** | **31** | **6** |
| 5.6 | MQR | SMA | 200.1±13.9 | 10 | 2 | 0.292±0.012 | 32 | 6 |
| 5.7 | YRR | SMA | 525.8±24.4 | 14 | 2 | 0.211±0.005 | 31 | 6 |
| 5.8 | FRR | SMA | 593.2±43.9 | 15 | 2 | 0.128±0.006 | 28 | 6 |
| 5.9 | RPK | SMA | −10.0±0.9 | 15 | 2 | 0.749±0.034 | 32 | 6 |
| 5.10 | ENP | SMA | −50.9±2.1 | 10 | 2 | 0.160±0.021 | 30 | 6 |
| **5.11** | **WRR** | **SMA** | **695.0±66.4** | **10** | **2** | **0.160±0.013** | **21** | **6** |
| 5.12 | LWI | SMA | 36.4±4.4 | 10 | 2 | 0.337±0.034 | 38 | 6 |
| 5.13 | MMA | SMA | 13.1±1.6 | 11 | 2 | 0.405±0.024 | 31 | 6 |
| 5.14 | FCK | SMA | 133.5±11.5 | 12 | 2 | 0.202±0.016 | 22 | 6 |
| 5.15 | WNK | SMA | 110.4±8.5 | 10 | 2 | 0.353±0.023 | 30 | 6 |
| 5.16 | LWR | SMA | 89.2±6.5 | 10 | 2 | 0.334±0.020 | 25 | 6 |
| 5.17 | ILR | SMA | 62.8±7.7 | 8 | 2 | 0.288±0.008 | 24 | 6 |
| 5.18 | LFR | SMA | 71.6±2.7 | 11 | 2 | 0.225±0.013 | 33 | 6 |
| 5.19 | YWR | SMA | 81.4±4.7 | 10 | 2 | 0.251±0.013 | 34 | 6 |
| 5.20 | FRK | SMA | 302.9±24.1 | 10 | 2 | 0.140±0.006 | 30 | 6 |
| 5.21 | MFI | SMA | 23.6±2.8 | 12 | 2 | 0.227±0.009 | 34 | 6 |
| **6.1** | **FAR** | **HVD** | **134.7±12.7** | **8** | **2** | **0.399±0.033** | **25** | **6** |
| 6.2 | FAR | QCP | 82.7±13.7 | 9 | 2 | 0.355±0.016 | 27 | 6 |
| 6.3 | FAR | QYK | 128.4±22.5 | 11 | 2 | 0.370±0.024 | 30 | 6 |
| 6.4 | FAR | HKP | 233.1±15.6 | 9 | 2 | 0.162±0.009 | 23 | 6 |
| 6.5 | FAR | QPS | 195.2±11.3 | 11 | 2 | 0.174±0.007 | 26 | 6 |
| 6.6 | FAR | SFA | 196.6±10.5 | 11 | 2 | 0.146±0.004 | 32 | 6 |
| 6.7 | FAR | HPS | 203.9±6.4 | 7 | 2 | 0.165±0.012 | 21 | 6 |
| **6.8** | **FAR** | **TIS** | **252.4±19.3** | **9** | **2** | **0.157±0.006** | **27** | **6** |
| 6.9 | FAR | MRP | 215.7±28.0 | 11 | 2 | 0.152±0.005 | 24 | 6 |
| 6.10 | FAR | HHY | 152.1±13.5 | 10 | 2 | 0.152±0.005 | 31 | 6 |
| 6.11 | FAR | VGI | 48.9±2.6 | 10 | 2 | 0.162±0.005 | 29 | 6 |
| 6.12 | FAR | KPS | 194.4±22.5 | 9 | 2 | 0.194±0.013 | 23 | 6 |
| 6.13 | FAR | FHR | 170.4±6.6 | 13 | 2 | 0.119±0.006 | 28 | 6 |
| 6.14 | FAR | SWG | 306.4±25.9 | 10 | 2 | 0.109±0.006 | 29 | 6 |
| 6.15 | FAR | TYY | −15.9±3.5 | 9 | 2 | 0.032±0.004 | 24 | 6 |

| | | | | | | | | |
|---|---|---|---|---|---|---|---|---|
| **6.16** | **FAR** | **YFL** | **560.0±69.0** | **6** | **2** | **0.087±0.007** | **21** | **6** |
| 6.17 | FAR | YFM | 615.8±75.5 | 8 | 2 | 0.069±0.005 | 24 | 6 |
| 6.18 | FAR | CWK | 335.1±30.6 | 9 | 2 | 0.090±0.003 | 28 | 6 |
| 6.19 | FAR | QWS | 213.3±13.1 | 7 | 2 | 0.098±0.005 | 24 | 6 |
| 6.20 | FAR | MKC | 203.9±13.5 | 12 | 2 | 0.158±0.004 | 28 | 6 |

**Supplementary Table S4**: Biosensor variants generated by combining left and right linker variants. The response ($\Delta F/F_0$) upon Fsk/IBMX stimulation and the basal brightness ($JF_{635}$/EGFP) are given. Additionally, number of cells and FOV are given. For $\Delta F/F_0$, FOV correspond to replicates; for basal brightness, six FOVs were measured from two replicates. The parental biosensor 3.5 = HaloAKAR1.0, as well as the selected biosensors from libraries L1 and L2, are given at the beginning of the table. The three selected biosensors are highlighted in bold. 7.6 = HaloAKAR2.0, 7.9 = HaloAKAR2.1, and 7.10 = HaloAKAR2.2.

| # | L1 | L2 | $\Delta F/F_0$±sem [%] | Cells | FOV | $JF_{635}$/EGFP±sem | Cells | FOV |
|---|---|---|---|---|---|---|---|---|
| 3.5 = HaloAKAR1.0 | FAR | SMA | 129.8±9.9 | 29 | 6 | 0.231±0.014 | 73 | 18 |
| 5.1 | QMR | SMA | 67.0±2.8 | 13 | 2 | 1.096±0.032 | 36 | 6 |
| 5.2 | CKK | SMA | 101.3±4.8 | 15 | 2 | 0.784±0.047 | 37 | 6 |
| 5.3 | MKR | SMA | 301.6±22.0 | 10 | 2 | 0.436±0.062 | 29 | 6 |
| 5.5 | MRR | SMA | 554.0±23.1 | 12 | 2 | 0.345±0.009 | 31 | 6 |
| 5.11 | WRR | SMA | 695.0±66.4 | 10 | 2 | 0.160±0.013 | 21 | 6 |
| 6.1 | FAR | HVD | 134.7±12.7 | 8 | 2 | 0.399±0.033 | 25 | 6 |
| 6.8 | FAR | TIS | 252.4±19.3 | 9 | 2 | 0.157±0.006 | 27 | 6 |
| 6.16 | FAR | YFL | 560.0±69.0 | 6 | 2 | 0.087±0.007 | 21 | 6 |
| 7.1 | QMR | HVD | 23.7±1.6 | 13 | 2 | 1.492±0.033 | 33 | 6 |
| 7.2 | CKK | HVD | 2.3±0.7 | 14 | 2 | 1.277±0.044 | 32 | 6 |
| 7.3 | MKR | HVD | 21.4±3.3 | 11 | 2 | 1.191±0.074 | 29 | 6 |
| 7.4 | MRR | HVD | 57.2±3.2 | 11 | 2 | 0.654±0.016 | 29 | 6 |
| 7.5 | WRR | HVD | 135.7±9.5 | 13 | 2 | 0.324±0.008 | 32 | 6 |
| **7.6 = HaloAKAR2.0** | **QMR** | **TIS** | **110.8±14.7** | **10** | **2** | **1.230±0.030** | **30** | **6** |
| 7.7 | CKK | TIS | 135.5±5.4 | 12 | 2 | 0.943±0.017 | 34 | 6 |
| 7.8 | MKR | TIS | 490.1±39.6 | 12 | 2 | 0.350±0.018 | 30 | 6 |
| **7.9 = HaloAKAR2.1** | **MRR** | **TIS** | **605.3±48.6** | **13** | **2** | **0.480±0.018** | **30** | **6** |
| **7.10 = HaloAKAR2.2** | **WRR** | **TIS** | **1346.0±79.0** | **12** | **2** | **0.211±0.008** | **30** | **6** |
| 7.11 | QMR | YFL | −25.16±1.4 | 10 | 2 | 0.615±0.040 | 29 | 6 |
| 7.12 | CKK | YFL | −32.7±2.6 | 7 | 2 | 0.168±0.025 | 22 | 6 |
| 7.13 | MKR | YFL | −18.03±1.7 | 7 | 2 | 0.106±0.004 | 30 | 6 |
| 7.14 | MRR | YFL | 3.1±0.5 | 9 | 2 | 0.152±0.012 | 25 | 6 |
| 7.15 | WRR | YFL | 71.5±12.6 | 9 | 2 | 0.177±0.013 | 31 | 6 |

**Supplementary Table S5**: Characterization of the three main biosensors HaloAKAR2.0, HaloAKAR2.1, and HaloAKAR2.2. The response ($\Delta F/F_0$) upon Fsk/IBMX stimulation, the basal brightness ($JF_{635}$/EGFP), and the maximum brightness compared to HaloTag7 ($JF_{635}$/EGFP) are given. Additionally, number of cells and FOV are given. For $\Delta F/F_0$, FOV corresponds to replicates; for basal brightness, six FOVs were measured from three replicates; and for activated brightness, eight FOVs from four replicates. The parental biosensor 3.5 = HaloAKAR1.0 is also given.

| # | L1 | L2 | $\Delta F/F_0$±sem [%] | Cells | FOV | Basal brightness $JF_{635}$/EGFP±sem | Cells | FOV | Max brightness $JF_{635}$/EGFP±sem | Cells | FOV |
|---|---|---|---|---|---|---|---|---|---|---|---|
| **HaloAKAR1.0** | FAR | SMA | 129.8±9.9 | 29 | 6 | 0.272±0.016 | 52 | 6 | - | - | - |
| **HaloAKAR2.0** | QMR | TIS | 104.4±5.1 | 24 | 3 | 1.323±0.048 | 54 | 6 | 0.299±0.007 | 58 | 8 |
| **HaloAKAR2.1** | MRR | TIS | 634±29 | 28 | 3 | 0.671±0.026 | 54 | 6 | 0.464±0.016 | 78 | 8 |
| **HaloAKAR2.2** | WRR | TIS | 1250±37 | 30 | 3 | 0.272±0.011 | 48 | 6 | 0.324±0.008 | 81 | 8 |
| **HaloTag7** | - | - | - | - | - | - | - | - | 2.488±0.033 | 75 | 8 |

**Supplementary Table S6**: Fit parameters from labeling experiments of biosensors and HaloTag with Alexa488-CA. sd = standard deviation.

| # | $k_{app}$±sd [$M^{-1}$ $s^{-1}$] | Literature[7] [$M^{-1}$ $s^{-1}$] |
|---|---|---|
| **HaloAKAR2.0** | $2.143 \pm 0.004 \cdot 10^3$ | - |
| **HaloAKAR2.1** | $0.8890 \pm 0.0007 \cdot 10^3$ | - |
| **HaloAKAR2.2** | $0.1865 \pm 0.0005 \cdot 10^3$ | - |
| **HaloTag7** | $25.15 \pm 0.07 \cdot 10^3$ | $25.7 \pm 0.1 \cdot 10^3$ |

**Supplementary Table S7**: Photophysical properties of HaloAKARs and HaloTag7 labeled with $JF_{635}$-CA. Absorbance maxima ($\lambda_{abs}$), excitation maxima ($\lambda_{ex}$), emission maxima ($\lambda_{em}$), extinction coefficients (ε) and quantum yields (ϕ) were measured in vitro, and phosphorylation was preformed via addition of $PKA_{cat}$ and ATP. Fluorescence lifetime (τ) was measured in living HeLa cells before and after stimulation with Fsk/IBMX. ND = not determined.

| # | Phosphorylation | $\lambda_{ex}/\lambda_{abs}$ [nm] | $\lambda_{em}$ [nm] | ε±sd [$M^{-1}$ $cm^{-1}$] | ϕ±sd | τ±sem [ns] | N [cells] |
|---|---|---|---|---|---|---|---|
| **HaloAKAR2.0** | - | 640/642 | 654 | 15,000±800 | 0.73±0.04 | 4.10±0.01 | 26 |
| | + | 640/642 | 654 | 21,300±1,100 | 0.74±0.08 | 4.19±0.01 | 26 |
| **HaloAKAR2.1** | - | 640/642 | 654 | 6,600±300 | 0.53±0.05 | 4.15±0.02 | 18 |
| | + | 640/642 | 654 | 17,500±1,800 | 0.78±0.04 | 4.23±0.1 | 25 |
| **HaloAKAR2.2** | - | 640/640 | 652 | 3,800±200 | 0.43±0.03 | 4.06±0.04 | 11 |
| | + | 640/642 | 652 | 16,200±900 | 0.84±0.02 | 4.22±0.1 | 24 |
| **HaloTag7**[1] | - | 640[1] | 656[1] | 81,000[1] | 0.75[1] | 3.84±0.01 | 18 |
| | “+” | ND | ND | ND | ND | 3.80±0.03 | 20 |

**Supplementary Table S8**: Survival fraction after 600 s bleaching.

| # | Survival fraction±sem [%] | Stimulated survival fraction±sem [%] |
|---|---|---|
| **HaloAKAR2.0** | 61.9±0.5 | 32.6±1.5 |
| **HaloAKAR2.1** | 71.2±2.3 | 14.5±0.8 |
| **HaloAKAR2.2** | 69.1±1.6 | 11.4±0.4 |
| **HaloCaMP1a** | 39.7±1.4 | 23.5±2.1 |
| **HaloTag7** | 75.5±0.6 | - |

### Supplementary Videos

**Supplementary Video S1:** PKA activity in isolated mouse islets upon acute glucose stimulation. Video of isolated mouse islets expressing HaloAKAR2.2-T2A-EGFP labeled with $JF_{635}$-CA responding to actuate stimulation with glucose (25 mM) at 2 min. Intensity increases corresponding to increased PKA activity can be seen at 5 min (3 min after addition). Scale bar, 50 µm.

### Methods

**General considerations:** HaloTag fluorophores ($JF_{635}$-CA, $JF_{585}$-CA, $JF_{526}$-CA, $JF_{639}$-CA, $JFX_{646}$-CA, and $JF_{669}$-CA) were kindly provided by L. Lavis (Janelia Research Campus) or purchased from commercial vendors (Alexa488-CA; Promega). Fluorophores were prepared as stock solutions in dry DMSO (Sigma) and diluted in the respective buffer such that the final concentration of DMSO did not exceed 1% v/v.
Drug treatments with forskolin 50 µM (Fsk, Calbiochem); 3-isobutyl-1methylxanthine 100 µM (IBMX, Sigma-Aldrich); H89 20 µM (Sigma); phorbol 12-myristate 13-acetate 100 ng $mL^{-1}$ (PMA, LC Laboratories); Gö6983 1 µM (Sigma); atrial natriuretic peptide 0.4 µM (ANP, AnaSpec); platelet-derived growth factor 50 ng $mL^{-1}$ (PDGF, Sigma-Aldrich); epidermal growth factor 100 ng $mL^{-1}$ (EGF, Sigma-Aldrich); tetraethylammonium chloride 20 mM (TEA, Sigma); ionomycin 1 µM (iono, Calbiochem); isoproterenol 1 µM (iso, Sigma); glucose 25 mM (Sigma) were performed as indicated.

**Plasmids:** A pET51b(+) vector (Novagen) was used for protein production in *E. coli* BL21. Proteins were N-terminally tagged with $His_{x10}$, followed by a TEV cleavage site (ENLYFQ|G). A pcDNA3.1 vector (Invitrogen) was used for transient expression in mammalian cells. HaloAKAR and its variants were fused to T2A-EGFP, CAAX, Lyn11, TOMM20, β-actin, CEP41, CLC, or an NES for expression in mammalian cells. Cloning was performed by Gibson assembly[8]. DNA was subsequently transferred to *E. coli* DH5α via heat-shock and plated on Lysogenic Broth (LB) agar plates with 100 µg $mL^{-1}$ ampicillin and incubated at 37 °C overnight. HaloAKAR-T/A variants, as well as PKC, Akt and ERK sensors, were cloned in a similar fashion.
For stable cell line generation, lentiviral plasmids were cloned via digestion of the pSIN-ELYK[9] plasmid with EcoRI and SpeI (ThermoFisher Scientific) and Gibson assembly[8] of a PCR-amplified insert.
pcDNA3-AKAR4 (Addgene plasmid # 61619)[5], pcDNA3-GR-AKAR3 (Addgene plasmid #173015)[6], pcDNA3.1(+)-ExRai-AKAR (Addgene plasmid # 118405)[3], pcDNA3.1(+)-ExRai-AKAR2 (Addgene plasmid # 161753)[4], pSIN-ELYK[9], pcDNA3-sapphireCKAR (Addgene plasmid # 118468)[3], pcDNA3-sapphireAKAR (Addgene plasmid # 118464)[3], pcDNA3-EKAR4 (Addgene plasmid # 174437)[10], pcDNA3-GFP11(x7)-Actin (Addgene plasmid # 181967)[11] were available in house and either used as entry plasmids or directly for transient transfection.
pcDNA5-H2B_Halo_T2A_EGFP, pET51b_rHCaMP, pcDNA5-CEP41_Halo, pcDNA5-TOM20_Halo_T2A_EGFP, pET51b-His-TEV-HaloTag7, pCDNA5/FRT/TO_HaloTag_T2A_EGFP were gifts from Kai Johnsson (Addgene plasmid # 135443[12], # 187106[2], # 135446[12], # 135443[12], # 167266[7], # 169327[13]); pAAV-snyapsin-HaloCaMP1a-EGFP, pAAV-snyapsin-HaloCaMP1b-EGFP, and pCAG-HASAP were gifts from Eric Schreiter (Addgene plasmid # 138327, # 138328, and # 138325)[1]; pEGFP-mNG2(11)_Clathrin light chain was a gift from Bo Huang (Addgene plasmid # 82608)[14]; RCaMP1d[15] was provided by L. Looger. These plasmids were used as template plasmids. CAAX (KKKKKSKTKCVIM), Lyn11 (MGCIKSKRKDKDP), and NES (MPPLERLTL) were cloned using overlapping primers.
psPAX2 and pMD2.G were gifts from Didier Trono (Addgene plasmid # 12260, # 12259) and directly used for lentiviral production.
CMV-B-GECO1 was a gift from Robert Campbell (Addgene plasmid # 32448)[16], Flamindo2, Pink Flamindo and Green cGull were gifts from Tetsuya Kitaguchi (Addgene plasmid # 73938[17], # 102356[18] and # 86867)[19], pCAG-G-Flamp1 was a gift from Jun Chu (Addgene plasmid # 188567)[20], and were directly used for transient transfection.

**Rational Engineering:** Initial HaloAKAR constructs were cloned in pcDNA3.1 starting from ExRaiAKAR1 (# 118405[3]), ExRaiAKAR2 (# 161753[4]), HaloCaMP1a (# 138327[1]), HaloCaMP1b (# 138328[1]), HASAP (# 138325[1]) and RHaloCaMP (# 187106[2]), according to Supplementary Figure S1 and Supplementary Table S1. Linkers were varied as indicated in Supplementary Table S1. All plasmids were cloned via Gibson cloning. The plasmids were subsequently transiently transfected into HeLa cells, and the biosensor responses tested upon Fsk/IBMX stimulation (see below). For selected variants of the rational screen, T2A-EGFP variants were cloned using Gibson cloning and the basal brightness ($JF_{635}$/EGFP) was determined (see below).

**Sort-Seq Screening:** Library screening was adapted from a previously published procedure[9]. The two libraries (L1 and L2) were generated using Golden Gate Assembly, varying the linker region preceding (L1) or following (L2, Supplementary Figure S2b) cpHaloTag. To this end, the receiver plasmid pSIN-PKA-substrate-Esp3I-LacZ-Esp3I-FHA1-T2A-EGFP was generated in two steps via PCR and Gibson cloning in pcDNA3.1, followed by subcloning of the whole cassette into the pSIN vector using restriction digestion (SpeI and EcoRI, ThermoFisher Scientific) and Gibson assembly.
To construct the biosensor, cpHaloTag was amplified by PCR with a forward primer containing an Esp3I site, an 8 bp connecting piece, the 9 bp (3 aa) long linker region, and the beginning of cpHaloTag; and a reverse primer containing an Esp3I site, a 7 bp connecting piece, the 9 bp (3 aa) long linker region, and the end of cpHaloTag. For L1, an NNK degenerate forward primer (Eton Bioscience) was used (NNKNNKNNK: $(4*2*2)^3$ = 32,768 DNA sequences) where all three amino acids (XXX) of the linker region preceding cpHaloTag were varied (XXX: $20^3$ = 8,000 protein variants). N represents an equimolar distribution of A, T, G, and C; K represents an equimolar distribution of T and G; and X represents any amino acid. For L2, an MNN degenerate reverse primer was used, randomizing all three amino acids following cpHaloTag. M represents an equimolar distribution of A and C. The sequences of the primers are listed in Supplementary Methods Table S1. The annealing temperature during PCR was varied from 55 to 72 °C. The resulting PCR fragments were purified via gel extraction and used to replace the LacZ domain in the receiver plasmid via Golden Gate assembly using Esp3I (NEB) and T4 ligase (Invitrogen). The products were purified and concentrated using DNA Cycle Pure Kit (Omega BioTec). The purified plasmids were transformed into ElectroMAX™ DH10B™ Cells (Invitrogen, >1 × $10^{10}$ CFU/µg) and grown in 250 mL LB cultures. The plasmids were purified using Qiagen HiSpeed Plasmid Maxi kit (Qiagen). The resulting plasmid libraries were validated using Sanger Sequencing (Genewiz).
*Lentivirus and stable cell line generation:* Next, the plasmid libraries were used to generate lentivirus in HEK293T cells. Briefly, a 10-cm dish of 60% confluent HEK293T cells were co-transfected with the pSin plasmid library (L1 or L2), the viral packaging plasmid psPAX2 (Addgene # 12260) and the envelop plasmid pMD2.G (Addgene # 12259) using Polyjet (SignaGen Laboratories). The medium was replaced with fresh medium 5 h after transfection. Viral particles were collected 48 h after transfection, filtered through a 0.45-µm filter (Cytiva), and concentrated using LentiX Concentrator (TakaRa). The virus titer was measured by flow cytometry in HeLa cells (SONY SH800). The plasmids of biosensor libraries were introduced into HeLa cells through viral infection with low multiplicity of infection (MOI = 0.1) allowing for a low copy number of plasmids per single cell: according to the Poisson distribution, less than 0.5% of cells should have two or more plasmid copies. Briefly, HeLa cells were seeded at 2 × $10^6$ cells in a 10-cm dish a day before viral transduction. Virus was added at an MOI of 0.1. Cells were sorted for EGFP-positive cells (ex: 488 nm, em: 525/50 nm) 72 h after transduction. The sorted cells were grown to confluency and

further used for library sorting. In addition to the cells for sorting, cells from the two libraries (L1 and L2) were kept as input control for sequencing, and cells were frozen for long-term storage.

*Library sorting:* One day before sorting, four 10-cm dishes ($4 \times 10^6$ HeLa cells each) were seeded for each library (2 dishes untreated, 2 dishes Fsk/IBMX stimulated). On the day of the sort, cells were labeled with $JF_{635}$-CA (500 nM for 3 h) and then washed twice with fresh medium. Two dishes per library were stimulated with Fsk/IBMX (50 µM and 100 µM) for 15 min. After stimulation, cells were washed with PBS, trypsinized, pelleted, resuspended in PBS + 10% FBS (+ Fsk/IBMX for the stimulated condition), passed through a cell strainer and kept on ice until sorting.

The expression-level-normalized intensity ($JF_{635}$/EGFP) was used to sort cells (EGFP ex: 488 nm, em: 525/50 nm; $JF_{635}$ = APC ex: 638 nm, em: 655/30 nm). Several controls were utilized to gate the desired cell populations. Plain HeLa cells were used to gate for live, singlet and fluorescent cells. HeLa cells expressing HaloAKAR1.0 labeled with $JF_{635}$-CA were used to gate for cells that express the sensor only (no EGFP). Unlabeled HeLa cells expressing HaloAKAR1.0-T2A-EGFP were used to gate for cells that show only EGFP fluorescence (unlabeled or non-expressers, Supplementary Figure S2c). HeLa cells expressing HaloAKAR1.0-T2A-EGFP labeled with $JF_{635}$-CA and stimulated with Fsk/IBMX were used to gate for high responders (high $JF_{635}$/EGFP ratio), and similarly the same cells untreated were used to gate for medium basal brightness (medium $JF_{635}$/EGFP ratio, Supplementary Figure S2d). After setting the gates, cells containing biosensor libraries were sorted into high $JF_{635}$/EGFP ratio and medium $JF_{635}$/EGFP ratio groups (Supplementary Figure S2). For each group, ~10% of biosensor-expressing cells were selected. This resulted in four groups: stimulated-high, stimulated-medium, untreated-high, and untreated-medium. FlowJo software 10 was used for data visualization (BD).

*Sequencing:* After sorting, total RNA was extracted from each pool of sorted cells (4 groups per library), as well as from the non-sorted input library, using Direct-zol RNA Microprep (Zymogen). Purified total RNA was used as a template for cDNA synthesis using SuperScript IV reverse transcriptase (Invitrogen) with a gene-specific primer (Supplementary Methods Table S2). Adaptor sequences with different indexes for Illumina sequencing (Supplementary Methods Table S2) were added to the prepared cDNA using PCR with <20 cycles. The PCR product was purified by gel-electrophoresis (2% agarose gel) via a Gel extraction Kit (Omega BIoTec). The purified amplicon libraries were then sequenced by Sanger sequencing (Genewiz) to verify the success of library preparation. High-throughput sequencing was performed on an Illumina NovaSeq 6000 at the UC San Diego IGM Genomics Center (paired-end 100 sequencing, >25 M reads per group).

*Analysis of the sequencing data:* FastQC[21] was used for quality control, and only data sets with quality >25 were used. Read 1 (R1) and read 2 (R2) data sets were analyzed independently. A custom bash script was used to identify sequences that contained the correct sequencing adaptors and the correct length of the linker region. The selected sequences were converted from nucleotide sequence to amino acid sequence and their occurrence counted. A custom R-script was then used to calculate the frequency of unique sequences ($f_v$) in counts per million (CPM) by normalizing the variant count in each group to the total number of sequencing reads. Next, enrichment ratios ($E_v$) were calculated, dividing $f_v$ of a sorted group by $f_{v,input}$ in the input library. Ideal biosensors are enriched ($E_v$ >1) in the stimulated-high ($E_{SH}$) and untreated-medium ($E_{UM}$) groups and not enriched ($E_v$ <1) in the stimulated-medium ($E_{SM}$) and untreated-high ($E_{UH}$, counter sort) groups. We selected biosensors fulfilling all four criteria and ranked them according to their product of $E_{SH}$ (stimulated-high) and $E_{UM}$ (untreated-medium), which was previously shown to correlate with biosensor performance[9]. Cut-offs were visually determined after plotting the ranked data (Supplementary Figure S2e) and correspond roughly to the top 20 candidates for each library. The selected biosensors were validated using fluorescence microscopy, measuring dynamic range and basal brightness (Table S3). Linkers from the L1 and L2 libraries leading to the highest performance were further combined and the resulting biosensors further validated (Table S4).

**Protein production and purification:** Proteins were expressed in *E. coli* strain BL21. LB cultures were grown at 37 °C to an optical density ($OD_{600nm}$) of 0.6, induced by the addition of isopropyl-β-D-thiogalactopyranoside (IPTG, 0.5 mM) and grown at 16 °C overnight. The cells were pelleted by centrifugation (3,100 g, 10 min, 4 °C), resuspended in lysis buffer (50 mM Tris, pH 7.4, 300 mM NaCl) containing Complete EDTA-free Protease Inhibitor Cocktail (Roche), and lysed by sonication (45 s sonication, 15 s rest, 7 min total, 70%, Sonics Vibra cells) on ice. The cell lysate was cleared by centrifugation (15,000 g, 20 min, 4 °C). Clarified lysate was loaded onto HisPur™ Ni-NTA resin (ThermoFisher Scientific). Unbound protein was washed away using wash buffer (50 mM Tris, pH 7.4, 300 mM NaCl, 10 mM imidazole), and bound protein was subsequently eluted using an imidazole gradient (10-300 mM) in 50 mM Tris, pH 7.4, 300 mM NaCl buffer. Eluted fractions were analyzed via SDS–PAGE (coomassie staining, see below), pooled and concentrated using Amicon Ultra-15 centrifugal filter device (10-kD cut-off, Millipore) followed by buffer exchange into 50 mM HEPES, 150 mM NaCl, pH 7.2 (<0.1 mM imidazole), leading to purity around 90% (verified by SDS-PAGE coomassie staining). The proteins were flash frozen and stored at −80 °C. Amino acid sequences are listed in the Supplementary Methods.

**Polyacrylamide gel electrophoresis (PAGE):** HaloAKAR proteins (9 µM, 20 µL) were labeled with $JFX_{646}$-CA (50 µM) in 50 mM HEPES, 150 mM NaCl, pH 7.2, for 1 h at room temperature. 4x Laemmli buffer (7 µL, BioRad) was added and the sample heated to 95 °C for 5 min. Proteins were separated by PAGE (4–20% 10 well Mini-Protean TGX, BioRad) and revealed by in-gel fluorescence using a ChemiDoc MP Imaging System (BioRad). $JFX_{646}$-CA-labeled proteins were imaged using epi-red illumination (ex: 635/25; em: 700/50 nm); total protein was visualized by coommassie staining (SimplyBlue™ SafeStain, Invitrogen) and colorimetric imaging.

**One-photon excitation and emission spectra:** Purified biosensor (2 µM) was incubated with $JF_{635}$-CA (1 µM) in modified kinase assay buffer (50 mM Tris-HCl, pH 7.5, 10 mM $MgCl_2$, 0.1 mM EDTA, 2 mM DTT) at room temperature for 24 h. After 24 h, excitation and emission spectra of the unphosphorylated biosensor were measured on a PTI QM-400 fluorimeter using FelixGX v.4.1.2 software (Horiba). Excitation scans were collected at 700 nm and excited from 540-680 nm, and emission scans were performed from 610-740 nm and excited at 590 nm. Then, PKAcat (0.1 µg $µL^{-1}$) and ATP (200 µM) were added to the samples and incubated for 30 min at 30 °C. Spectra of the phosphorylated biosensor were then measured.

**Extinction coefficient measurements:** Purified biosensor or HaloTag7 (20 µM) was incubated with $JF_{635}$-CA (10 µM) in modified kinase assay buffer (50 mM Tris-HCl, pH 7.5, 10 mM $MgCl_2$, 0.1 mM EDTA, 2 mM DTT) at room temperature for 24 h. After 24 h, absorbance spectra of the unphosphorylated biosensor were measured on a combined fluorescence and absorbance spectrometer (Duetta Horiba). Then, PKAcat (0.2 µg $µL^{-1}$) and ATP (400 µM) were added to the samples and incubated for 45 min at 30 °C. Spectra of the phosphorylated biosensor were then measured. Data were baseline corrected, and the maximum absorbance values at 640 nm were used to calculate the extinction coefficient relative to the reported value for HaloTag7-$JF_{635}$ (81,000 $M^{-1}$ $cm^{-1}$)[1].

**Relative quantum yields:** In addition to measuring the absorbance spectra (see extinction coefficient measurements), emission spectra (610-750 nm) excited at 590 nm were measured before and after phosphorylation on a combined fluorescence and absorbance spectrometer (Duetta Horiba). All spectra were baseline corrected and the area below the emission peak calculated. The calculated extinction coefficients (ε) and measured areas (A) were used to calculate the relative quantum yields (ϕ) from the reported value for HaloTag7-$JF_{635}$ (0.75)[1] according to equation (1).

$$\varphi_{sample} = \frac{A_{sample}}{A_{HT}} \frac{\varepsilon_{HT}}{\varepsilon_{sample}} \varphi_{HT} \quad (1)$$

**Kinetics by plate reader:** Labeling kinetics of HaloTag7 and HaloAKARs were measured by time-course fluorescence polarization measurements using Alexa488-CA (Promega) as previously described[7]. Briefly, fluorescence polarization was measured on a CLARIOstar microplate reader (BMG Labtech) in black nonbinding flat-bottom 96-well plates (Corning Inc.) with a final volume of 200 µL at 37 °C. Excitation was performed at 482/16 nm, emission was collected at 530/40 nm, gains and measurement height were optimized from reference measurements (free fluorophore in buffer and fully labeled HaloTag7 in buffer). Labeling reactions were started by injecting Alexa488-CA (final concentration 50 nM) into wells containing different protein concentrations (3.2-25.6 µM) in 50 mM HEPES, 150 mM NaCl, 0.5 g $L^{-1}$ BSA, pH 7.2, using the built-in injector module. A baseline was measured for free Alexa488-CA without any protein. All measurements were performed in triplicate.

The recorded data were fit using a kinetic model (2) using DynaFit software[22] as previously described. The Y-axis offset was fixed using the average fluorescence polarization measured for free Alexa488-CA, as was the fluorophore substrate concentration (S = 50 nM).

$$P + S \xrightarrow{k_{app}} PS \quad (2)$$

**Cell culture and transfection:** HeLa cells were cultured in Dulbecco's modified Eagle medium (DMEM, Gibco) containing 1 g $L^{-1}$ glucose, 10% (v/v) fetal bovine serum (FBS, Sigma) and 1% (v/v) penicillin-streptomycin (Pen-Strep, Sigma-Aldrich). HEK293T were cultured in DMEM (Gibco) containing 4.5 g $L^{-1}$ glucose, 10% (v/v) FBS (Sigma) and 1% (v/v) Pen-Strep (Sigma-Aldrich). NIH3T3 cells were cultured in DMEM (Gibco) containing 1 g $L^{-1}$ glucose and supplemented with 10% (v/v) fetal calf serum and 1% (v/v) Pen-Strep (Sigma-Aldrich). MIN6 pancreatic β-cells were cultured in DMEM (Gibco) containing 4.5 g $L^{-1}$ glucose, supplemented with 10% (v/v) FBS, 1% (v/v) Pen-Strep and 50 µM β-mercaptoethanol. All cells were maintained in a humidified incubator at 37 °C with a 5% $CO_2$ atmosphere. Cells were split every 2-3 days or at confluency. Cell lines were regularly tested for mycoplasma contamination via DNA staining.

For imaging experiments, cells were seeded onto sterile quartered 35-mm glass-bottomed dishes (Cellvis) and grown to 50–70% confluence. For HEK293T cells, dishes were poly-*D*-lysine (ThermoFisher) coated before seeding. Transient transfection of HeLa, HEK293T and MIN6 cells was performed using Lipofectamine 2000 (Invitrogen) according to the manufacturer's recommendations: DNA (125 ng) was mixed with OptiMEM (13 µL, Gibco), and Lipofectamine 2000 (0.4 µL) was mixed with OptiMEM (13 µL). The solutions were incubated for 5 min at room temperature, then mixed and incubated for 10 min. The prepared DNA-Lipofectamine complex was added to one of the wells and incubated for 16 h, after which medium was changed to fresh medium. HeLa and HEK293T cells were imaged 20-24 h and MIN6 48 h after transfection. Transfection for three- or four-color imaging was also performed using Lipofectamine 2000

(Invitrogen) using a total of 375 ng (3 x 125 ng) and 500 ng (4 x 125 ng) DNA, respectively. Transfection for five-color imaging was performed using calcium phosphate precipitation. Briefly, 625 ng of DNA (5 x 125 ng) were mixed with $CaCl_2$ (2 µL, 2 M) and $H_2O$ (22.5 µL). The DNA mixture was added dropwise to HBS (25 µL; 50 mM HEPES, 0.28 M NaCl, 1.5 mM $Na_2HPO_4$, pH 7.0) while vortexing. The resulting mixture was incubated for 10 min and added to the cells. NIH3T3 cells were changed to serum-free DMEM immediately before transfection using Polyjet (SignaGen). Briefly, DNA (250 ng) was mixed with serum-free DMEM (13 µL), and Polyjet (6 µL) was mixed with serum-free DMEM (13 µL). The solutions were incubated for 5 min at room temperature, then mixed and incubated for 10 min. The transfection complex was added to the wells and incubated for 6 h. Medium was changed to fresh serum-free DMEM and starvation was continued for another 18 h.

**Stable cell lines**: HEK293T stably expressing HaloAKAR2.1-NES-T2A-EGFP-CAAX were generated via lentivirus. First, pSin-HaloAKAR2.1-NES-T2A-EGFP-KRAS was used to generate lentivirus in HEK293T cells. Briefly, a 10-cm dish of 60% confluent HEK293T cells were co-transfected with the pSin plasmid, the viral packaging plasmid psPAX2 (Addgene # 12260) and the envelop plasmid pMD2.G (Addgene # 12259) using Polyjet (SignaGen Laboratories). After 24 h transfection, the medium was replaced with fresh medium. Viral particles were collected 48 h after transfection and filtered through a 0.45 µm filter (Cytiva). Lentivirus was then added to HEK293T cells at 50% confluency. 72 h after transduction, medium containing puromycin (1 µg $mL^{-1}$) was added to select for biosensor-expressing cells. Cells were grown to confluency, and expression was verified by fluorescence microscopy.

**Spheroid generation:** HEK293T cells stably expressing HAKAR2.1-NES-T2A-EGFP-CAAX were grown into spheroids over the course of 4 d. Glass-bottom imaging dishes were coated with a thin layer of Matrigel (Corning), then cured at 37 °C for 30 min. Quartered 35-mm glass-bottom dishes (Cellvis) were used to grow spheroids for spinning-disc confocal microscopy, 35 mm glass-bottom dishes (Cellvis) were used to grow spheroids for 2P fluorescence microscopy, and 25-mm circular #1 microscope cover glasses (Fisher Scientific) were used to grow spheroids for lattice light-sheet microscopy. Cells were trypsinized and seeded on top of the cured Matrigel at a density of $7 \times 10^3$ cells per dish, then grown in DMEM (Gibco) containing 4.5 g $L^{-1}$ glucose, 10% (v/v) FBS (Sigma) and 1% (v/v) Pen-Strep (Sigma-Aldrich) supplemented with 2% of Matrigel for 4 d.

**Islet sample preparation:** Male C57BL/6J mice (Jackson Laboratory; RRID: IMSR_JAX:000664, lean, 10 weeks old) were maintained at 22 °C on a 12 h light/12 h dark cycle. After mice were euthanized, pancreatic islets were isolated by injecting collagenase P digestion solution (1.4 mg $mL^{-1}$; Roche) into the pancreas via the common bile duct. The perfused pancreas was digested at 37 °C and repeatedly vortexed until homogenously dissolved. Digestion was quenched by ice-cold complete medium (RPMI, 10 % FBS). After centrifugation (3 min, 300 g, 4 °C), the supernatant was aspirated and the cell pellet washed twice with cold complete medium and filtered through a 500-µm strainer (#U-CMN-500-B, Component Supply). The filtered solution was spun down and resuspended in a smaller volume to slowly place the islets on top of a 70-µm cell strainer (#352350, Corning). The strainer containing the islets was inverted into a 6-cm tissue culture dish (#83.3901.500, SARSTEDT) containing 5 mL complete RPMI medium. Islets were handpicked under a stereoscopic microscope and seeded into quartered 35-mm glass-bottomed dishes in RPMI 1640 medium with 10% (v/v) FBS, 1% (v/v) Pen-Strep, and 11.1 mM glucose and allowed to recover for 24 h. The next day, $2.5 \times 10^{10}$ vg $mL^{-1}$ AAV8-Ins-HAKAR2.1-T2A-EGFP with an inducible insulin promotor[23]

(VectorBuilder) were added to the islets and incubated for 6-7 d. On the day of imaging, islets were labeled with $JF_{635}$-CA (500 nM) in islet imaging buffer at low glucose (10 mM HEPES, pH 7.4, 1.2 mM $KH_2PO_4$, 2.6 mM $CaCl_2$, 4.7 mM KCl, 1.2 mM $MgSO_4$, 115 mM NaCl, 0.5% BSA, 2.8 mM glucose) for 3 h, then washed twice in the same buffer and imaged as described below. Glucose stimulation was performed to a final concentration of 25 mM glucose.
All animal procedures were approved by the local Animal Care and Use Committee and performed in accordance with the University of California, San Diego Research Guidelines for the Care and Use of Laboratory Animals.

**Labeling and sample preparation for live-cell imaging:** Cells were labeled with the indicated fluorophores (fluorophore-CA, 500 nM for 3 h) at 37 °C in corresponding cell culture medium, washed twice with the same medium (1 min each, 37 °C) and changed into Hank's balanced salt solution (HBSS, Gibco). For cAMP, cGMP, PKA, PKC, and $Ca^{2+}$ imaging in HeLa, HEK293T, and MIN6, cells were imaged immediately. For ERK imaging in HEK293T, cells were additionally starved in HBSS for 30 min prior to imaging. For Akt imaging in NIH3T3, labeling (fluorophore-CA, 500 nM for 3 h) was performed in serum-free DMEM medium, followed by washes in the same medium (twice for 1 min, 37 °C). Cells were then changed into HBSS and imaged after 30 min.

**Widefield microscopy:** Images were acquired on a Zeiss AxioObserver Z1 microscope (Carl Zeiss) equipped with a 40×/1.3 NA objective, a Photometrics Evolve 512 EMCCD (Photometrics) camera and a Lambda 10-2 filter-changer (Sutter Instruments) controlled by METAFLUOR 7.10 software (Molecular Devices). Exposure times ranged between 50 and 500 ms, with the EM gain set from 10 to 50. An OD filter of OD = 0.6 was used unless otherwise stated, and images were acquired every 30 s unless otherwise stated.
*HaloAKAR measurements:* Far-red intensity was imaged using an ET640/30 excitation filter, a T660lpxr dichroic mirror, and an ET700/75 emission filter (far-red filter set); GFP intensity using an HQ480/30 excitation filter, a 505dcxr dichroic mirror, and an HQ535/45 emission filter (GFP filter set). For basal brightness measurements, far-red and GFP images were acquired before drug addition.
*Brightness comparison with HT7:* For brightness comparison of the activated biosensor, far-red and GFP images were acquired 3 min after Fsk/IBMX stimulation of HeLa cells expressing HaloAKAR2.x-T2A-EGFP or HaloTag7-T2A-EGFP labeled with $JF_{635}$-CA as described above.
*Bleaching experiments:* For bleaching experiments of cells expressing HaloAKAR2.x-T2A-EGFP, HaloCaMP1a-EGFP, or HaloTag7-T2A-EGFP, far-red intensity was measured every 1 s using the far-red filter set. To achieve efficient bleaching, the OD filter was removed. Stimulated cells were treated with Fsk/IBMX 3 min prior to image acquisition. For the measurements of HaloTag7 itself, cells were labeled with diluted $JF_{635}$-CA (500 pM for 3 h) to not saturate the detector.
*HaloAKAR labeled with different fluorophores:* $JF_{635}$-CA, $JF_{639}$-CA, $JFX_{646}$-CA, and $JF_{669}$-CA labeled cells were imaged with the far-red filter set. $JF_{585}$-CA labeled cells were imaged using an ET555/25 excitation filter, a ZT568rdc dichroic mirror, and an ET605/52 emission filter (RFP filter set). $JF_{526}$-CA labeled cells were imaged using an HQ495/10 excitation filter, a 515dcxr dichroic mirror, and an HQ535/25 emission filter (YFP filter set).
*Dose-response experiments:* HeLa cells expressing HaloAKAR2.2-EGFP labeled with $JF_{635}$-CA or ExRaiAKAR2 were imaged using the far-red filter set or dual GFP excitation-ratio imaging using an HQ480/30 excitation filter, 505dcxr dichroic mirror, and ET405/40 excitation filter, 505dcxr dichroic mirror, and HQ535/45

emission filter. Indicated doses of Fsk were added, and a maximum dose of Fsk/IBMX was added after 8 min to measure maximum biosensor response.

*Biosensor multiplexing:* Three-color multiplexing: G-Flamp-1 intensity was imaged using the GFP filter set; RCaMP1d intensity was imaged using the RFP filter set; Flamindo2 intensity was imaged using the YFP filter set; HaloAKAR labeled with $JF_{635}$-CA was imaged with the far-red filter set.

Four-color multiplexing: SapphireAKAR intensity was imaged using a ET380/10 excitation filter, 505dcrx dichroic mirror, and 535/45 emission filter (T-sapphire filter set); Flamindo2 intensity was imaged using the YFP filter set; pinkFlamindo intensity was imaged using the RFP filter set; HaloAKAR labeled with $JF_{635}$-CA was imaged with the far-red filter set.

Five-color multiplexing: BGeco intensity was imaged using a ET380/10 excitation filter, 450dcxru dichroic mirror, and HQ475/40 emission filter (BFP filter set); SapphireCKAR intensity was imaged using the T-sapphire filter set; cGull intensity was imaged using the YFP filter set; pinkFlamindo intensity was imaged using the RFP filter set; HaloAKAR labeled with $JF_{635}$-CA was imaged with the far-red filter set. Due to the low intensity of SapphireCKAR, the OD filter was removed.

*General image analysis:* Raw fluorescence images were corrected by subtracting the background fluorescence intensity of a cell-free region from the emission intensities of biosensor-expressing cells. The resulting time courses were normalized by dividing the intensity at each time point by the value at time zero ($F/F_0$), which was defined as the time point at the beginning of the experiment. For ExRai-AKAR2, GFP excitation ratios (F480nm/F405nm) were calculated at each time point and normalized to time point zero ($R/R_0$). Maximum intensity ($\Delta F/F_0$) changes were calculated as $(F_{max} - F_0)/F_0$, where $F_{max}$ is the maximum value recorded after stimulation and $F_0$ is the initial intensity at the beginning of the experiment. Brightness comparisons were calculated by dividing the background-subtracted fluorescence intensities of the far-red ($JF_{635}$) by the green (EGFP) channel. For dose-response analysis, ($\Delta F/F_0$) or ($\Delta R/R_0$) after 8 min of Fsk treatment was divided by ($\Delta F/F_0$) or ($\Delta R/R_0$) after 5 min of Fsk/IBMX treatment. Graphs were plotted using GraphPad Prism 8 (GraphPad Software).

**Confocal microscopy:** HEK293T spheroid samples stably expressing HAKAR2.1-NES-T2A-EGFP-CAAX were grown on Matrigel (Corning)-coated quartered 35-mm glass-bottom dishes (Cellvis) as described above. Spheroids were labeled with $JF_{635}$-CA (500 nM for 3 h) at 37 °C in HEK293T cell culture medium, washed with HEK293T cell culture medium, and placed in HBSS for imaging. Images were acquired on an Eclipse Ti2-E microscope (Nikon) equipped with a Plan Apo Lambda 40x/0.95 NA air objective (Nikon), a solid-state laser light engine with 405 nm, 446 nm, 477 nm, 520 nm, 546 nm, 638 nm, and 749 nm laser lines (Lumencor Celesta), an ORCA-Fusion digital CMOS camera (Hamamatsu), and an X-Light V3 spinning-disc with 50-µm pinholes (CrestOptics) controlled by NIS-Elements software (version 5.21.01, High content analysis package, Nikon). Far-red and EGFP intensities were imaged using triggered acquisition of the 638 nm and 477 nm laser lines, with a 440/40 nm, 520/20 nm, 607/34 nm, 694/35 nm, and 809/81 nm quad dichroic emission filter.

*4D imaging:* 16-bit z-stack images (18 slices) were collected at intervals of 3 µm throughout the volume of the spheroid. Both far-red (150 ms) and EGFP (150 ms) channels were measured every 30 s for 20 min. Spheroids were stimulated with 50 µM Fsk/IBMX at t = 2 min and 20 µM H89 at t = 10 min.

*Median plane:* 16-bit images were collected in the far-red (200 ms) and EGFP (200 ms) channels every 30 s for 20 min. Spheroids were stimulated with 50 µM Fsk/IBMX at t = 2 min and 20 µM H89 at t = 10 min.

*Image analysis:* One representative z-plane of the EGFP channel was registered using the ImageJ Plugin "Register Virtual Stack Slices" using a rigid feature model and rigid (translate + rotate) registration model.

The same transform was then applied to the same z-plane of the far-red channel using the “transform Virtual Stack Slices” Plugin[24]. The first frame of the registered EGFP time course was used to segment cells using Cellpose v2.0[25,26]. Background-subtracted far-red intensity values in the selected z-plane were measured at each time point, averaged within each segmented cell, and normalized to the average far-red intensity for each cell at t = 0 min.

**Lattice light-sheet microscopy:** HEK293T spheroids stably expressing HAKAR2.1-NES-T2A-EGFP-CAAX were grown on Matrigel-coated 25 mm #1 coverslips (Fisher Scientific) as described above. Spheroids were labeled with $JF_{635}$-CA (500 nM for 3 h) at 37 °C in HEK293T cell culture medium and washed with phenol-red-free DMEM (Gibco) containing 4.5 g $L^{-1}$ glucose, 10% (v/v) FBS (Sigma) and 1% (v/v) Pen-Strep (Sigma-Aldrich). Spheroids were then imaged on a custom-built lattice light-sheet microscope designed by the Betzig Lab (HHMI Janelia/UC Berkeley)[27,28] in the same medium under 5% $CO_2$ and at 37 °C. The microscope design is similar to that described in[27], and differences are described in[28]. In addition, the system uses Hamamatsu Orca Fusion BT sCMOS cameras. 488 nm and 640 nm lasers were used to excite EGFP and HAKAR2.1-$JF_{635}$, respectively. 50 ms exposure time was used and a ET525/50m-2p8 filter (Chroma) for EGFP and a BLP01-647R filter (Semrock) for $JF_{635}$. A Multiple Bessel Beam Light-Sheet Pattern with $NA_{Max}$ = 0.4, $NA_{Min}$ = 0.38 was used, with a sheet length of 75 µm. Measured resolution in x-y and z were 330 nm and 700 nm, respectively. The volume size was 2730 x 1548 x 546 pixels, or 300.30 x 170.28 x 60.06 µm, with isotropic pixel size 110 nm after coverglass transformation. Spheroids were measured before and after addition of Fsk/IBMX.
*Image analysis:* To perform 3D cell segmentation, we used custom Python code. Briefly, the raw EGFP channel (plasma membrane) was first preprocessed using sample-stage deconvolution, deskewing, and rotation to form the “membrane” z-stack. The membrane z-stack was then thresholded, dilated, and inverted to generate the seed labels for the 3D watershed segmentation. The watershed segmentation used the seed labels to flood the “membrane” z-stack to create the 3D masks for each individual cell in the spheroid. Cell masks were visually inspected, and inaccurate masks removed. The cell boundary showed negligible movement between measurements. Thus, the segmented cell masks generated from the first frame were applied to the second frame.
To analyze the fluorescence intensity change of HaloAKAR-$JF_{635}$, the raw HaloAKAR-$JF_{635}$ channel was preprocessed using deskewing and rotation (deconvolution was skipped to avoid intensity manipulations). The segmented cell masks prepared above were then utilized to crop the HaloAKAR-$JF_{635}$ z-stacks at the two time points. Specifically, the average fluorescence intensity within a cell mask was background corrected, normalized to timepoint t = 0 min, and taken as the normalized fluorescence intensity of that particular cell ($F/F_0$) and used to calculate $\Delta F/F_0$. Imaris (Oxford Instruments) was used to generate 3D visualizations.

**2P microscopy**: HEK293T spheroid samples stably expressing HAKAR2.1-NES-T2A-EGFP-CAAX were grown on Matrigel-coated 35-mm glass-bottom dishes (Cellvis) as described above. Spheroids were labeled with $JF_{635}$-CA or $JF_{585}$-CA (500 nM for 3 h) at 37 °C in HEK293T cell culture medium and washed with phenol-red-free DMEM (Gibco) containing 4.5 g $L^{-1}$ glucose, 10% (v/v) FBS (Sigma) and 1% (v/v) Pen-Strep (Sigma-Aldrich). Spheroids were imaged on an upright laser-scanning microscope (DIY multiphoton, Olympus) equipped with a 25x water objective (XLPLN, WMP2, 1.05 NA, Olympus). A tunable picosecond laser (PicoEmerald, APE) was utilized to perform 2P excitation at 1031 nm. Back-scattered fluorescence emission was collected using a 495-540 BP, 575-645 BP and 570 DC

(Olympus) filter set. The detected signal was then fed into the FV-OSR software module (Olympus) integrated with the FV3000 microscope system to generate images during laser scanning. Before drug treatment, a single image of size 512 × 512 pixels was acquired with a dwell time of 40 µs and an imaging speed of approximately 10 s per image. After drug treatment, a series of images (time-course) was collected every 30 s for 10 cycles. The measured z-plane images were analyzed identically to the spinning disc confocal data (see above).

**STED microscopy:** Imaging was performed on a Leica Stellaris 8 (Leica Microsystems) equipped with a Leica scanhead, a white-light laser (WLL) for excitation, a 775 nm depletion laser, Leica HyDS and HyDX detectors, and an HC PL APO 86x1.20 motCORR STED water immersion objective. Excitation was performed at 630 nm, and emission was collected between 650-750 nm, and temporal gating was performed from 0.6 – 8 ns. Pixel size was 35.3 nm (640x640 pixels), pixel dwell time 6.15 µs, and 4-line accumulations were collected for all images. Laser powers were kept constant, so images were comparable (WLL at 80% overall, using 30% for excitation; 775 nm laser at 100% overall, using 50% for depletion). The objective was equipped with a motorized correction collar (motCORR, Leica Microsystems), and its position was optimized for every sample to maximize signal-to-noise ratio. Alignment and performance of the STED microscope were checked every day using fluorescent beads as described in detail in[29]. Briefly, 20 nm diameter fluorescent microspheres (F8782 Thermo Fisher) were immobilized on poly-*D*-lysine (A3890401 Thermo Fisher)-coated #1.5 glass cover slips (Zeiss), mounted on microscope slides (VWR), sealed with nail polish, and stored at 4 °C for less than 1 month. After automatic alignment of the Stellaris 8 in LAS-X (Leica Microsystems), expert alignment was performed to ensure perfect spatial overlap of the excitation and depletion beam using the fluorescent bead sample under the same imaging parameters as described above. Confocal and STED images of HaloAKAR2.1-$JF_{635}$ in HeLa cells were acquired line-wise interleaved before and 2 min (Fsk/IBMX or no-treatment) or 3 min (Fsk) after drug treatment.

*Diameter analysis:* For the quantification of the clathrin-coated pit diameter, the perpendicular line profile of a 105 nm (3 pixels)-wide section was measured in ImageJ/Fiji[30,31]. The normalized profile was fitted to a Gaussian function (10) and the FWHM derived thereof (11). Multiple line profiles were measured within individual cells imaged on different days.

$$y = y_0 + \frac{A}{w\sqrt{\frac{\pi}{2}}} e^{-2\frac{(x-x_c)^2}{w^2}} \tag{10}$$

$$FWHM = w\sqrt{2\ln(2)} \tag{11}$$

*Photobleaching measurements:* 30 consecutive STED frames (12.9 s per frame) were acquired under the same imaging settings as described above, and the background-corrected intensity within a ROI was compared over time.

*Pit-level PKA activity:* Custom Matlab (MathWorks) code (including the Image Processing Toolbox) was used to segment clathrin-coated pits before and after drug stimulation. Briefly, the raw image undergoes pre-processing with difference-of-Gaussian filtering to enhance spot detection (kernel size = 5; sigma1 = 5; sigma2 = 10) and in a second step noise reduction via a Gaussian blur with *imgaussfilt* (sigma = 2). After binarization with Otsu's method, objects are filtered for circularity (0.2 < x < 1) and area (25 pixels < x) and labeled with a unique identifier. The Euclidean distance between the labeled objects detected in the image before and after drug stimulation was calculated using *dist2*. The pairings with the shortest distances were kept, and additional pairings with distances longer than 14 pixels were removed. We then calculated the mean $F/F_0$ and $\Delta F/F_0$ for the paired pits after background-correcting the images.

**Scanning confocal microscopy for FLIM microscopy and islet experiments:** *FLIM:* Measurements were performed on a Leica SP8 FALCON microscope (Leica Microsystems) equipped with a Leica TCS SP8 X scanhead; a SuperK white-light laser, Leica HyD SMD detectors, and a HC PL APO CS2 40x1.10 water immersion objective. Excitation was performed at 633 nm at a pulse frequency of 80 MHz and a dwell time of 3.2 µs. Emission was collected at 645-750 nm. To determine the fluorescence lifetime, cells expressing HaloTag7 or cytosolic HaloAKAR variants were imaged, collecting maximum 1,000 photons per pixel (512x512). The acquired cell images were intensity thresholded to remove background signal, and single-cell ROIs were drawn. Fluorescence lifetimes were calculated in the LAS X software (Leica Microsystems) by fitting mono-exponential decay models (n-exponential reconvolution) to the decay ($X^2 < 1.5$). Three replicates were measured before and after Fsk/IBMX addition.
*Islet experiments:* Islets prepared as described above were imaged on the same microscope using simultaneous excitation at 488 nm and 633 nm and a dwell time of 1.6 µs. Dual emission was collected at 500-580 nm and 650-750 nm. Images were acquired every 30 s (1024x1024) for a total of 20 min. Images were background subtracted, and ROIs were drawn manually in ImageJ/Fiji[30,31]. Mean fluorescence intensity was extracted for each ROI at each time point and normalized to t = 0 min. Normalized fluorescence intensity traces were then plotted over time.

**Software and image processing:** Statistical analysis was performed using GraphPad Prism 10 (GraphPad Software). Microsoft Excel and R[32], including packages readxl[33] and tidyverse[34], were used to process data. Images were processed with METAFLUOR 7.10, Cellpose v2.0[25,26], ImageJ/Fiji[30,31] and macros written therein unless otherwise stated.

**Statistics and Reproducibility:** Representative microscopy images and time courses are shown. The number of replicates is indicated with the experiments. Replicates stem from distinct samples unless otherwise stated. Details regarding statistical testing are given with the experiments in the Supplementary Figures.

**Protein sequences**

blue: Hisx10-tag, green: TEV cleavage site, orange: linker region,

HaloAKAR2.0

MHHHHHHHHHHENLYFQGLRRATLVDGGTGGSELSQMRETFQAFRTTDVGRKLIIDQNVFIEGTLPMGVVRPLTEVE
MDHYREPFLNPVDREPLWRFPNELPIAGEPANIVALVEEYMDWLHQSPVPKLLFWGTPGVLIPPAEAARLAKSLPNCK
AVDIGPGLNLLQEDNPDLIGSEIARWLSTLEISGEPTTGGSGGTGGSGGTGGSMAEIGTGFPFDPHYVEVLGERMHYVD
VGPRDGTPVLFLHGNPTSSYVWRNIIPHVAPTHRCIAPDLIGMGKSDKPDLGYFFDDHVRFMDAFIEALGLEEVVLVIH
DWGSALGFHWAKRNPERVKGIAFMEFIRPIPTWDEWTISKFSQEQIGENIVCRVICTTGQIPIRDLSADISQVLKEKRSIK
KVWTFGRNPACDYHLGNISRLSNKHFQILLGEDGNLLLNDISTNGTWLNGQKVEKNSNQLLSQGDEITVGVGVESDILS
LVIFINDKFKQCLEQNKVDR

HaloAKAR2.1

MHHHHHHHHHHENLYFQGLRRATLVDGGTGGSELSMRRETFQAFRTTDVGRKLIIDQNVFIEGTLPMGVVRPLTEVE
MDHYREPFLNPVDREPLWRFPNELPIAGEPANIVALVEEYMDWLHQSPVPKLLFWGTPGVLIPPAEAARLAKSLPNCK
AVDIGPGLNLLQEDNPDLIGSEIARWLSTLEISGEPTTGGSGGTGGSGGTGGSMAEIGTGFPFDPHYVEVLGERMHYVD
VGPRDGTPVLFLHGNPTSSYVWRNIIPHVAPTHRCIAPDLIGMGKSDKPDLGYFFDDHVRFMDAFIEALGLEEVVLVIH
DWGSALGFHWAKRNPERVKGIAFMEFIRPIPTWDEWTISKFSQEQIGENIVCRVICTTGQIPIRDLSADISQVLKEKRSIK
KVWTFGRNPACDYHLGNISRLSNKHFQILLGEDGNLLLNDISTNGTWLNGQKVEKNSNQLLSQGDEITVGVGVESDILS
LVIFINDKFKQCLEQNKVDR

HaloAKAR2.2

MHHHHHHHHHHENLYFQGLRRATLVDGGTGGSELSWRRETFQAFRTTDVGRKLIIDQNVFIEGTLPMGVVRPLTEVE
MDHYREPFLNPVDREPLWRFPNELPIAGEPANIVALVEEYMDWLHQSPVPKLLFWGTPGVLIPPAEAARLAKSLPNCK
AVDIGPGLNLLQEDNPDLIGSEIARWLSTLEISGEPTTGGSGGTGGSGGTGGSMAEIGTGFPFDPHYVEVLGERMHYVD
VGPRDGTPVLFLHGNPTSSYVWRNIIPHVAPTHRCIAPDLIGMGKSDKPDLGYFFDDHVRFMDAFIEALGLEEVVLVIH
DWGSALGFHWAKRNPERVKGIAFMEFIRPIPTWDEWTISKFSQEQIGENIVCRVICTTGQIPIRDLSADISQVLKEKRSIK
KVWTFGRNPACDYHLGNISRLSNKHFQILLGEDGNLLLNDISTNGTWLNGQKVEKNSNQLLSQGDEITVGVGVESDILS
LVIFINDKFKQCLEQNKVDR

**Supplementary Methods Table 1**: Primer sequences used to generate libraries L1 and L2.

| # | FP | RP |
|---|---|---|
| **L1** | CACCACACGTCTCGAGCGAGCTCAGC**NNKNNKNNK** GAGACCTTCCAGGCCTTCCG | CACCACACGTCTCGTGAGAAAACTTAGCCATGCTCCAT TCGTCCCAGGTCGGGATAG |
| **L2** | CACCACACGTCTCGAGCGAGCTCAGCTTTGCCCGCG AGACCTTCCAGGCCTTCCG | CACCACACGTCTCGTGAGAAAACTT**MNNMNNMNN**C CATTCGTCCCAGGTCGGGATAG |

**Supplementary Methods Table 2**: Primer sequences used to generate amplicon libraries. Green: p5 sequence, red: sequencing primer Rd1, orange: annealing region plasmid specific, yellow: p7 sequence, black: i7 index unique to each group, purple: sequencing primer Rd2.

| Name | Sequence 5’-3’ |
|---|---|
| cDNA primer | CCTAAATGATAGTCACAGGCTGG |
| **L1-FP** | AATGATACGGCGACCACCGAGATCTACACTCTTTCCCTACACGACGCTCTTCCGATCT GTCGCGCCACCCTGGTG |
| **L2-FP** | AATGATACGGCGACCACCGAGATCTACACTCTTTCCCTACACGACGCTCTTCCGATCT CGCGTCAAAGGTATTGCATTTATGG |
| **L1-input-RP** | CAAGCAGAAGACGGCATACGAGATCGTGATGTGACTGGAGTTCAGACGTGTGCTCT TCCGATCCCATCGGCAGCGTACCCTCG |
| **L1-stim-high-RP** | CAAGCAGAAGACGGCATACGAGATACATCGGTGACTGGAGTTCAGACGTGTGCTCT TCCGATCCCATCGGCAGCGTACCCTCG |
| **L1-stim-med-RP** | CAAGCAGAAGACGGCATACGAGATGCCTAAGTGACTGGAGTTCAGACGTGTGCTCT TCCGATCCCATCGGCAGCGTACCCTCG |
| **L1-untreat-high-RP** | CAAGCAGAAGACGGCATACGAGATTGGTCAGTGACTGGAGTTCAGACGTGTGCTCT TCCGATCCCATCGGCAGCGTACCCTCG |
| **L1-untreat-med-RP** | CAAGCAGAAGACGGCATACGAGATCACTGTGTGACTGGAGTTCAGACGTGTGCTCT TCCGATCCCATCGGCAGCGTACCCTCG |
| **L2-input-RP** | CAAGCAGAAGACGGCATACGAGATATTGGCGTGACTGGAGTTCAGACGTGTGCTCT TCCGATCGGAATTTGACCCGTGGTACAAATG |
| **L2-stim-high-RP** | CAAGCAGAAGACGGCATACGAGATGATCTGGTGACTGGAGTTCAGACGTGTGCTCT TCCGATCGGAATTTGACCCGTGGTACAAATG |
| **L2-stim-med-RP** | CAAGCAGAAGACGGCATACGAGATTCAAGTGTGACTGGAGTTCAGACGTGTGCTCT TCCGATCGGAATTTGACCCGTGGTACAAATG |
| **L2-untreat-high-RP** | CAAGCAGAAGACGGCATACGAGATCTGATCGTGACTGGAGTTCAGACGTGTGCTCT TCCGATCGGAATTTGACCCGTGGTACAAATG |
| **L2-untreat-med-RP** | CAAGCAGAAGACGGCATACGAGATAAGCTAGTGACTGGAGTTCAGACGTGTGCTCT TCCGATCGGAATTTGACCCGTGGTACAAATG |
